# Golgi-IP, a novel tool for multimodal analysis of Golgi molecular content

**DOI:** 10.1101/2022.11.22.517583

**Authors:** Rotimi Fasimoye, Wentao Dong, Raja S. Nirujogi, Eshaan S. Rawat, Miharu Iguchi, Kwamina Nyame, Toan K. Phung, Enrico Bagnoli, Alan Prescott, Dario R. Alessi, Monther Abu-Remaileh

**Affiliations:** Medical Research Council (MRC) Protein Phosphorylation and Ubiquitylation Unit, School of Life Sciences, University of Dundee, Dow Street, Dundee DD1 5EH, U.K.; Aligning Science Across Parkinson’s (ASAP) Collaborative Research Network, Chevy Chase, MD, 20815, USA; Department of Chemical Engineering, Stanford University, Stanford, CA 94305, USA; Department of Genetics, Stanford University, Stanford, CA 94305, USA; The Institute for Chemistry, Engineering & Medicine for Human Health (ChEM-H), Stanford University, Stanford, CA 94305, USA; Department of Biochemistry, Stanford University School of Medicine, Stanford, CA 94305, USA; Dundee Imaging Facility, School of Life Sciences, University of Dundee, Dow Street, Dundee DD1 5EH, U.K.

**Keywords:** Golgi-IP, Golgi apparatus, proteomics, metabolomics, lipidomics, uridine-diphosphate-hexose, uridine-diphosphate-N-acetyl-hexosamine, SLC35A2

## Abstract

The Golgi is a membrane-bound organelle that is essential for protein and lipid biosynthesis. It represents a central trafficking hub that sorts proteins and lipids to various destinations or for secretion from the cell. The Golgi has emerged as a docking platform for cellular signalling pathways including LRRK2 kinase whose deregulation leads to Parkinson disease. Golgi dysfunction is associated with a broad spectrum of diseases including cancer, neurodegeneration, and cardiovascular diseases. To allow the study of the Golgi at high resolution, we report a rapid immunoprecipitation technique (Golgi-IP) to isolate intact Golgi mini-stacks for subsequent analysis of their content. By fusing the Golgi resident protein TMEM115 to three tandem HA epitopes (GolgiTAG), we purified the Golgi using Golgi-IP with minimal contamination from other compartments. We then established an analysis pipeline using liquid chromatography coupled with mass spectrometry to characterize the human Golgi proteome, metabolome and lipidome. Subcellular proteomics confirmed known Golgi proteins and identified novel ones. Metabolite profiling established the first known human Golgi metabolome and revealed the selective enrichment of uridine-diphosphate (UDP) sugars and their derivatives, which is consistent with their roles in protein and lipid glycosylation. Furthermore, targeted metabolomics validated SLC35A2 as the subcellular transporter for UDP-hexose. Finally, lipidomics analysis showed that phospholipids including phosphatidylcholine, phosphatidylinositol and phosphatidylserine are the most abundant Golgi lipids and that glycosphingolipids are enriched in this compartment. Altogether, our work establishes a comprehensive molecular map of the human Golgi and provides a powerful method to study the Golgi with high precision in health and disease states.

**Significance:** The Golgi is central to protein and lipid processing. It senses and responds to diverse cell states to allow trafficking of macromolecules based on cellular demands. Traditional techniques for purifying the Golgi shaped our understanding of its functions, however such methods are too slow to preserve the labile Golgi metabolome and transient protein interactions. Here, we overcome this issue through the development of a method for the rapid capture of intact Golgi from human cells using organelle-specific immunoprecipitation (Golgi-IP). Using high resolution mass spectrometry, we demonstrate that our approach allows the unbiased characterization of the Golgi proteome, metabolome and lipidome. Thus, we believe that the Golgi-IP will be useful for the study of the Golgi in health and disease states.

## Introduction

The Golgi is a membrane-bound organelle central to protein and lipid processing, sorting and secretion. After synthesis in the endoplasmic reticulum (ER), secreted and organellar proteins undergo post-translational modifications in the Golgi including glycosylation, phosphorylation and tyrosine sulfation, processes that are essential for structural diversification, functional maturation and signaling (1, 2). Golgi dysfunction is associated with a multitude of human disorders including cancer, neurodegeneration and cardiovascular diseases (3–5).

To perform its functions, the Golgi relies on an adequate supply of metabolites required for modifying proteins and lipids (6, 7). Furthermore, the emerging dynamic nature of the Golgi suggests that it is capable of sensing and responding to diverse cell states to allow trafficking of macromolecules based on cellular demands (8, 9).

While traditional techniques for purifying the Golgi from mammalian cells, such as density-based centrifugation, helped shape our current understanding of its cellular roles, such methods are too slow to preserve what is likely a labile Golgi metabolome and transient protein interactions that regulate its functions. To overcome this, we used insights from our work and other’s to purify cellular organelles (10–18) to develop a novel method for the rapid capture of intact Golgi from human cells using organelle-specific immunoprecipitation (Golgi-IP).

Using our Golgi-IP method, we present for the first time a comprehensive characterization of the human Golgi proteome, metabolome and lipidome. Golgi-IP successfully enriched previously reported Golgi proteins while revealing novel proteins associated with this organelle. Furthermore, uridine-diphosphate (UDP) sugars and their derivatives as well as phospholipids were enriched in our untargeted metabolomics and lipidomics analyses, respectively. Finally, we used the Golgi-IP method to show that loss of the UDP-galactose transporter SLC35A2 (19, 20) leads to depletion of UDP-hexose specifically in the Golgi while its levels in the whole cell remain unchanged.

These data show that the Golgi-IP method coupled with modern omics technologies provides a powerful approach to study, in an unbiased manner, the molecular content of the Golgi and to uncover its regulation and function in health and disease states.

## Results

### Development and validation of the Golgi-IP method

Rapid purification of cellular organelles including mitochondria (12, 14, 15, 17), lysosomes (10, 11) peroxisomes (16) and early-endosomes (13) have allowed the analysis of their molecular content at unprecedented resolution. To develop an analogous approach to isolate intact Golgi, we generated a fusion protein comprised of the Golgi-specific transmembrane protein, TMEM115 (21, 22) whose N- and C-termini face the cytoplasm, fused to three tandem HA-epitopes at its C-terminus (TMEM115-3xHA; referred to as GolgiTAG) (Fig. 1A). Although the exact function of TMEM115 is not known, it interacts with the Conserved Oligomeric Golgi (COG) complex that is essential for the localization of Golgi resident enzymes and the tethering of transport vesicles to the Golgi (23). Endogenous TMEM115 has an estimated copy number of ~80,000 in HEK293 cells (See Methods).

**Figure 1:**
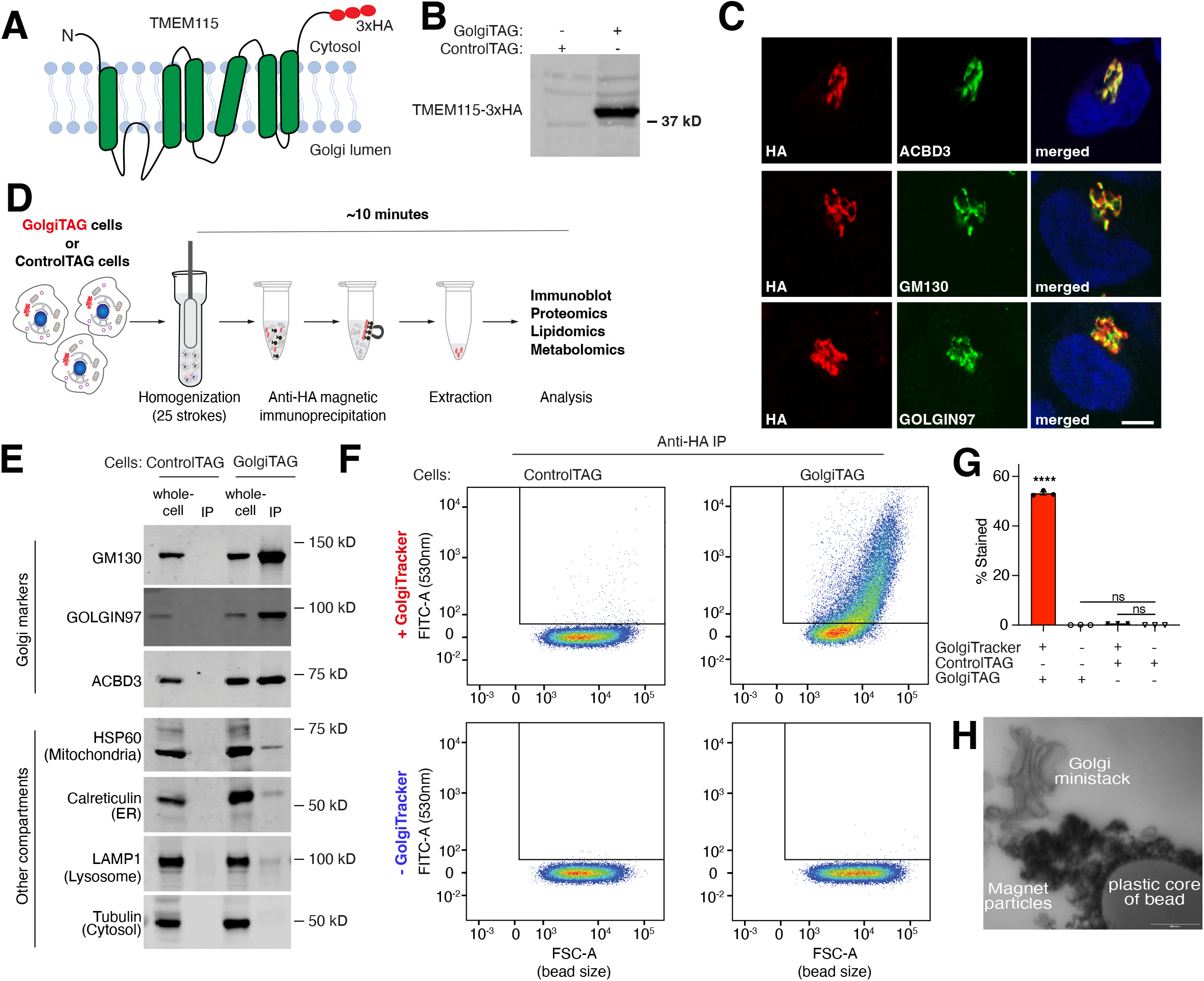
Development of the Golgi immunoprecipitation (Golgi-IP) method. (A) A depiction of the domain structure of the Golgi transmembrane protein TMEM115 fused to 3x-HA tag. (B) Immunoblot analysis of HEK293 cells stably expressing the TMEM115-3x-HA tag (GolgiTAG) using the HA antibody. (C) Immunofluorescence analyses confirm the Golgi localization of the GolgiTAG. HA in red channel and Golgi markers in green channel. The right panel displays a merged green and red channel. Scale bar is 1 μm. (D) Representation of the workflow of the Golgi-IP method (see methods section and protocols.io (dx.doi.org/10.17504/protocols.io.6qpvrdjrogmk/v1 for details). (E) Immunoblot analyses confirm that Golgi-IP isolates pure Golgi. Whole-cell lysates (2 μg) as well as the resuspended immunoprecipitates (IPs) (2 μg) were subjected to immunoblotting with the indicated compartment-specific antibodies. (F) Golgi-IP enriches for intact Golgi. Control HEK293 and HEK293 cells stably expressing the GolgiTAG were stained with and without 5 μM BODIPY™ FL C5-Ceramide (N-(4,4-Difluoro-5,7-Dimethyl-4-Bora-3a,4a-Diaza-s-Indacene-3-Pentanoyl)Sphingosine) (GolgiTracker) for 30 minutes at 37°C. Cells were lysed and subjected to Golgi-IP as in (E) except that the IP fraction was resuspended in KPBS and subjected to flow cytometry analysis using the FITC 530 nm channel. The beads bound to Golgi were first gated and selected as described in Fig. S1A. The selected beads were then analysed based on emission intensity in the FITC 530 nm (Y-axis) and bead size (Forward side scatter FSC, X-axis). (G) Quantitation of the percentage of beads stained with the GolgiTracker of 3 replicates. Data presented as mean ± SEM (One-way ANOVA). (H) Golgi ministacks are purified by Golgi-IP. As in (E) except that resuspended IPs from GolgiTAG cells were analyzed using Transmission Electron Microscopy. Scale bar 200 nm.

To immunopurify the Golgi from human cells, we generated human embryonic kidney cells (HEK293) stably expressing the GolgiTAG (Fig. 1A, 1B). As a control, we generated HEK293 cells stably expressing the empty backbone vector used to express the GolgiTAG, which retains the region encoding for the puromycin resistance and the 3xHA peptide (ControlTAG). Immunofluorescence microscopy confirmed that the GolgiTAG co-localizes with markers of the Golgi cisternae (ACBD3), *cis*-Golgi network (GM130/GOLGA2) and the *trans*-Golgi network (Golgin97/GOLGA1) (Fig. 1C). Of importance, the expression of the tag did not cause Golgi fragmentation (Fig. 1C), which is commonly observed upon depletion or over-expression of other Golgi proteins (24). After validating the localization of the GolgiTAG, we next homogenized GolgiTAG and ControlTAG HEK293 cells in isotonic potassium supplemented phosphate-buffer saline (KPBS) and performed an anti-HA immunoprecipitation of the Golgi using magnetic beads (Fig. 1D and methods). To minimize the leakage of metabolites and the degradation of labile content, the Golgi-IP was optimized to take ~10 minutes to isolate Golgi, starting with live cells. Immunoblot analyses of the bead-bound fraction revealed that GM130, GOLGIN97 and ACBD3 Golgi proteins were enriched in the immunoprecipitates (IPs) from GolgiTAG cells compared to markers for ER (Calreticulin), mitochondria (HSP60), cytosol (Tubulin) and lysosomes (LAMP1), which were depleted (Fig. 1E). No enrichment was observed in IPs from ControlTAG cells (Fig. 1E). Furthermore, to test if Golgi-IP preserves the metabolite content of purified organelles, cells were pre-treated with the Golgi-specific ceramide-based fluorescent dye (GolgiTracker, see methods) before Golgi-IP. Indeed, IPs from GolgiTAG cells were enriched for GolgiTracker, while no signal was observed in those from ControlTAG cells (Fig. 1F, 1G, Fig. S1A). Finally, transmission electron micrographs of the bound fraction showed that our preparation contained largely intact Golgi ministacks (Fig.1H), further verifying the integrity of the immunoprecipitated Golgi.

### Golgi-IP proteomics defines the Golgi proteome and reveals novel Golgi-associated proteins

To determine the proteomic landscape of the human Golgi, we used high- resolution data-independent acquisition (DIA) liquid chromatography/ tandem mass spectrometry (LC/MS-MS), to analyze the protein content of whole-cell and IP lysates derived from GolgiTAG- and ControlTAG cells (Fig. 2A, 2B, Fig. S1B-S1G, Table S1, and see methods). As expected, principal component and correlation matrix analyses revealed the distinct proteomic profile of IPs derived from GolgiTAG cells when compared to those derived from ControlTAG cells or the whole-cell fractions of either line (Fig. 2B). Note that no major change in global protein abundances was observed between lysates collected from GolgiTAG and ControlTAG whole cells (Fig. S2A, Table S2), thus the expression of TMEM115-3xHA has no detrimental effects on the cellular proteome. To further analyze this dataset, we first curated a comprehensive list of 1,363 proteins that were previously annotated as Golgi proteins in the compartment database or recently published sub-cellular organelle proteomic studies (25–30) (Table S3). In this curated list we included the estimated protein copy numbers for each protein in HEK293 whole-cell extracts (Table S3). These range from high abundance ~10 million copies per cell (TAGLN2, PARK7, CALU and RAB1A) to low abundance <1000 copies per cell (MACF1, COPA, and TMED5) (Table S3, Fig. S2B). Of the ~7000 proteins detected in our dataset, 527 were significantly at least twice as abundant in the isolated GolgiTAG-IP compared to ControlTAG-IP (adjusted p-value with 1% permutation-based FDR). 360 out of the 527 proteins were previously reported as Golgi-associated proteins in our curated list whereas the remaining 167 proteins were not (Fig. 2C and Table S1). Furthermore, 463 proteins were deemed Golgi-associated proteins when enrichment was calculated by comparing protein abundance in GolgiTAG-IP to that of the whole-cell (at least 2-fold enrichment, adjusted p-value with 1% permutation-based FDR) (Fig. 2D, Table S1). Among the identified proteins are the well-characterized Golgi proteins ACBD3 (5.3-fold) GM130/GOLGA2 (7.8-fold), Golgin-97/GOLGA1 (7.5-fold), B3GALT6 (13.1-fold), the Golgi luminal protein SDF4 (11.2-fold), GOLGA4 (4.9-fold), Rab29 (1.8-fold), Rab1B (2.7-fold), MAN2A1 (8.9-fold) and MAN2A2 (10.9-fold) (Fig. 2E and Table S1). Furthermore, we analyzed the GolgiTAG isolated proteins (GolgiTAG-IP/Control-IP and GolgiTAG-IP/whole-cell) for machineries required for Golgi functions including glycosylation, protein phosphorylation and dephosphorylation and protein ubiquitylation (Fig. S2C, S2D). As expected, the vast majority of the glycosylation-related proteins are indeed detected and enriched in the GolgiTAG IPs (Table S1).

**Figure 2:**
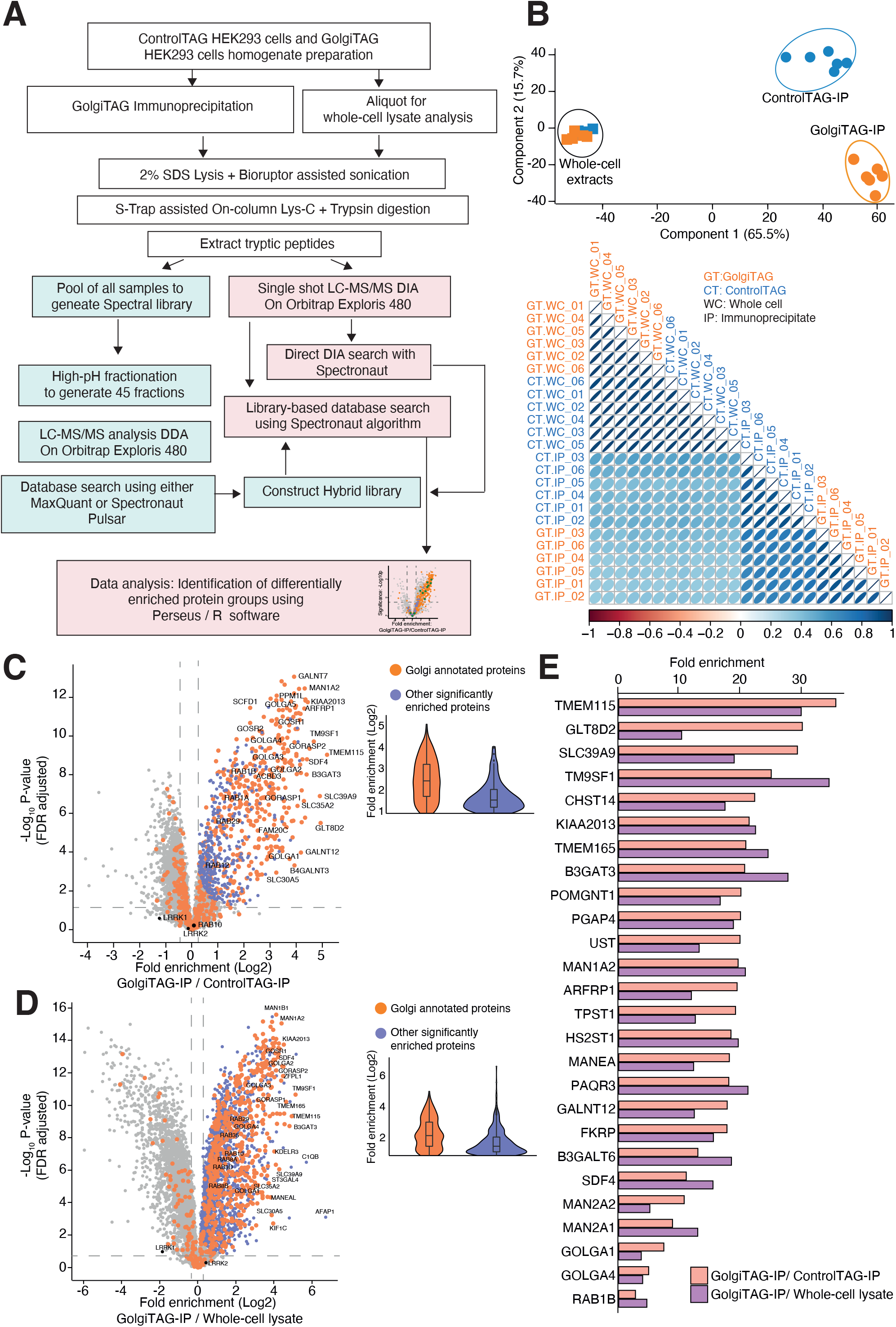
Quantitative proteomic analysis of Golgi derived from HEK293 cells. (A) Depiction of the workflow of the Data Independent Acquisition (DIA)-based mass spectrometry analysis of whole-cell lysates and HA immunoprecipitates (IPs) from Control and GolgiTAG expressing HEK293 cells. (B) Principal component analysis of DIA mass spectrometry data of whole-cell lysates (squares) and IPs (circle) from the Control (blue) and GolgiTAG expressing HEK293 cells (Orange) (upper panel). Lower panel present a global Pearson correlation of proteomics data of all samples. The control immunoblots for this experiment are presented in Fig. S1B. (C) Volcano plot of the fold enrichment of proteins between IPs from GolgiTAG and Control HEK293 cells (p-value adjusted for 1% Permutation-based FDR correction, s0=0.1) (left panel). The orange dots indicate known Golgi annotated proteins curated from databases described in (Table S3). Purple dots represent proteins enriched in the GolgiTAG samples that are not annotated as Golgi localized proteins. Selected well-studied Golgi proteins are denoted with gene names. The adjacent violin plots (right panel) show the significantly enriched proteins in GolgiTAG over Control IPs. (D) As in (C) except that the fold enrichment in protein abundances was calculated by comparing the IP fraction and the whole-cell lysate of GolgiTAG cells. (E) Bar graph presentation of the relative fold enrichment of top 15 proteins enriched in the IP fraction from GolgiTAG cells compared to those from ControlTAG cells (orange) or to whole-cell lysate from GolgiTAG cells (purple).

From the list of 360 proteins identified as enriched in the Golgi, we performed immunofluorescence analysis of the localization of SLC39A9 (29.7-fold) (31), TM9SF1 (25.4-fold) (32) and KIAA2013 (21.4-fold) (27–29) and confirmed previous reports that these proteins co-localize with the *trans*-Golgi network protein, GCC185 (33, 34) (Fig 3A). As there was no previous immunofluorescence data for KIAA2013, we treated cells with nocodazole to induce microtubule depolymerization, a well-established approach to verify Golgi compartment localization (33, 35). We found that nocodazole treatment does not disrupt the co-localization of KIAA2013 with GCC185 (Fig. 3A), further confirming KIAA2013 as a Golgi resident protein.

**Figure 3:**
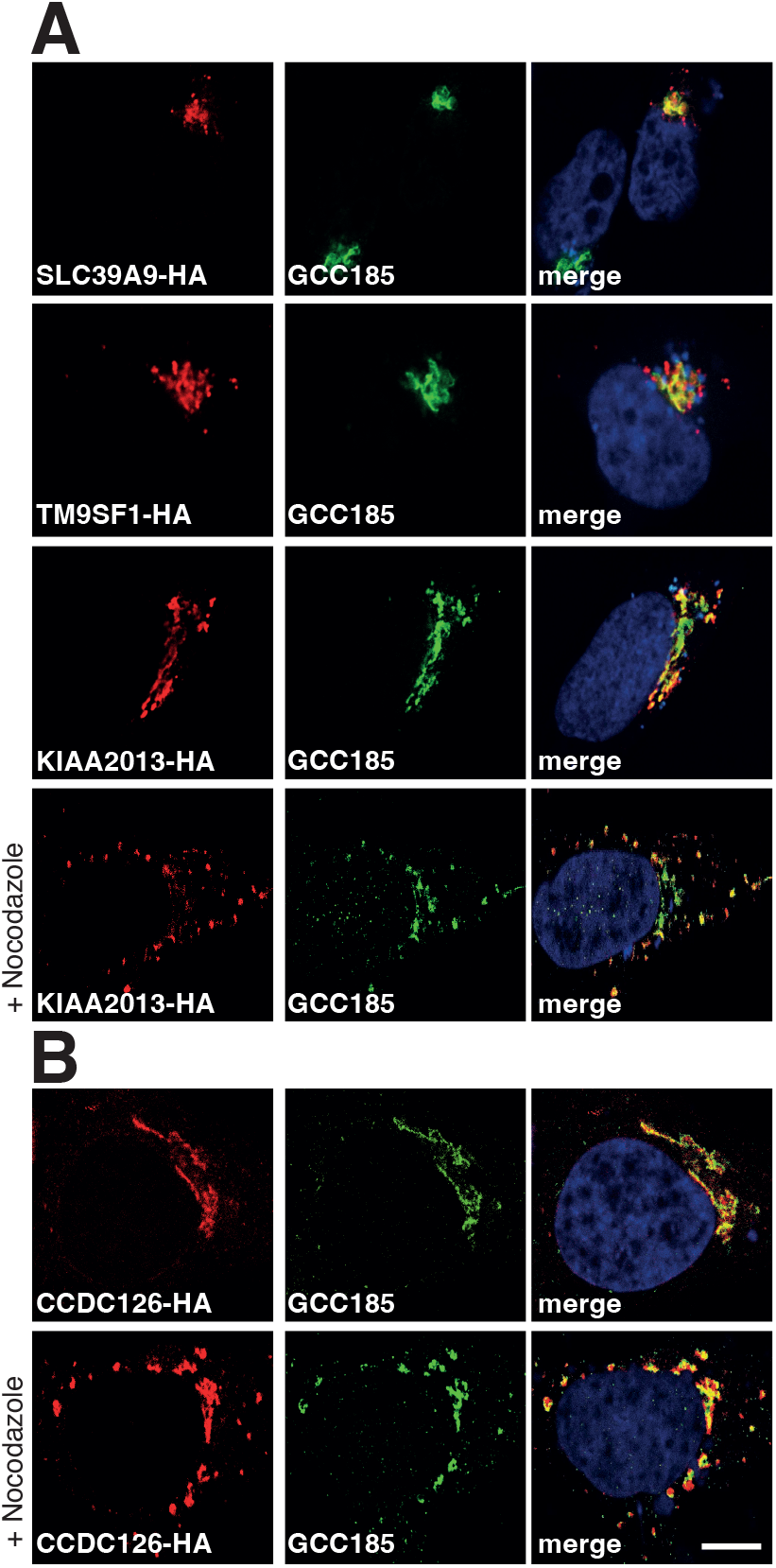
Immunofluorescence of proteins enriched in the Golgi. (A and B) Validation of Golgi localization of proteins found in the GolgiTAG IP. HeLa cells were transiently transfected with plasmids encoding the indicated proteins fused to a C-terminal HA tag. Cells were then fixed and stained with rat anti-HA (red, left panels) and rabbit anti-GCC185 (green, middle panels). The Right panel displays the merged red and green channel. Nuclei were stained with DAPI (blue). For nocodazole treated samples, cells were treated with ± 30 μM nocodazole for 1 h at 37°C 24 h post transfection. Scale bar is 1 μm.

To study potential novel Golgi-associated proteins, we turned to the list of 167 proteins that show significant enrichment (≥ 2-fold and adjusted p-value with 1% FDR) in our GolgiTAG-IPs but have not been reported as potential Golgi proteins (Table S3 and Table S1). By manually searching the Uniprot database, we identified 42 proteins to be annotated as putative ER proteins and 55 proteins that are associated with endosomes, lysosomes and centrioles. Of the remaining 70 proteins, we tested a few for their potential Golgi localization using over expression coupled with immunofluorescence analyses; these included DIPK1A (17.5-fold), C5orf22 (16.7-fold), CCDC126 (11.2-fold), TMEM219 (11.0-fold) and ABHD13 (10.2-fold). Of these, we observed a strong co-localization of CCDC126 with GCC185 that was not impacted by nocodazole treatment (Fig. 3B). In contrast, we didn’t observe strong co-localization of DIPK1A, C5orf22, TMEM219 ABHD13 with GCC185 (Fig. S3).

### The metabolic landscape of the human Golgi

Rapid organelle purification provides an unprecedented opportunity to analyze the labile metabolite content of cellular compartments (10, 11, 16, 17, 36). To determine the metabolite content of the Golgi, we performed untargeted metabolomics using LC/MS on IPs from GolgiTAG or ControlTAG cells. Consistent with their role in the Golgi (37), we found, using unbiased analysis, uridine-diphosphate (UDP) species to be among the most enriched metabolites in the Golgi (Fig. 4A and Table S4). Indeed, targeted analyses confirmed that UDP-hexose-uronate, UDP-N-acetyl-hexosamine and UDP-hexose were enriched 20.9-fold,10.9-fold, and 9.2-fold in IPs from GolgiTAG cells when compared to those from ControlTAG cells, respectively (Fig. 4B). Importantly, whole-cell levels of these species were similar between GolgiTAG and ControlTAG cells (Fig. 4B). In contrast to UDP-containing species, we did not observe any enrichment of the abundant cytosolic metabolite lactate in the Golgi samples (Fig. 4C), indicating that our method isolates pure Golgi.

**Figure 4:**
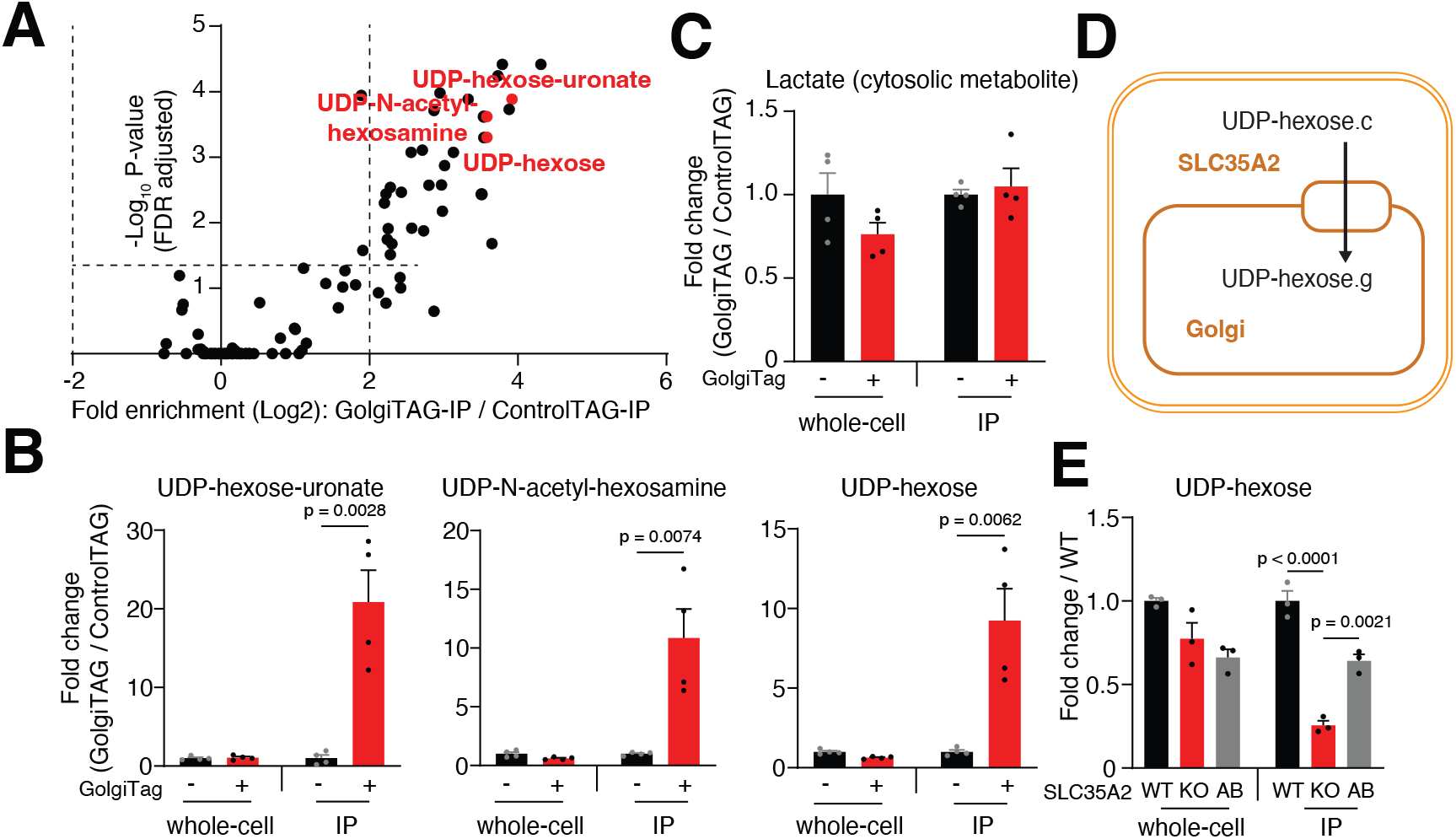
Golgi-IP metabolomics identified UDP-sugars as the main Golgi metabolites and pinpointed substrates for subcellular transporters. (A) UDP-hexose-uronate, UDP-N-acetyl-hexosamine and UDP-hexose are the most enriched species in the Golgi. Volcano plot depicting the fold enrichment of metabolites between IPs derived from Control and GolgiTAG HEK293 cells (p-value corrected by the Benjamini-Hochberg method, FDR=5%, *n=* 4). Data available in Table S4 (B) UDP-hexose-uronate, UDP-N-acetyl-hexosamine and UDP-hexose are enriched only in the Golgi while the wholecell level is not affected by the GolgiTAG. Targeted analyses showing fold change in the abundance of UDP species in the IP and whole-cell fractions of GolgiTAG compared to those of ControlTAG cells. (C) Lactate, an example of an abundant cytosolic metabolite, is not enriched in GolgiTAG IPs. (D) The transporter function of SLC35A2. SLC35A2 transfers UDP-hexose from the cytoplasm to the Golgi. g indicates the Golgi pool and c indicates the cytosolic pool. (E) Targeted analysis showing the fold change in the abundance of UDP-hexose in the IP and whole-cell fractions of *SLC35A2* KO and add-back (AB) HEK293 cells compared to those of wildtype (WT) cells. Data are presented as mean ± SEM (n= 4) for A-C and ± SEM (n= 3) for E. Statistical tests: two-tailed unpaired t-test for B-C and one-way ANOVA with Tukey’s HSD post-hoc for E.

In addition to UDP-sugars, we identified several metabolites in the Golgi (Table S4), including phosphocreatine and glutathione (Fig. S4). Phosphocreatine has been implicated in high-energy Golgi signalling involving coupling of creatine kinase and *cis*-Golgi matrix protein GM130 (38). In addition, glutathione is believed to be critical in maintaining Golgi oxidative homeostasis (39), and its depletion has been shown to compromise post-Golgi trafficking (40). Another class of metabolites found to be enriched in the Golgi were those related to phosphatidylcholine (PC): phosphocholine and glycerophosphocholine (GPC) (Fig. S4). As discussed later, PC is the most enriched lipid class in the Golgi. PC is synthesized *de novo* via the CDP-choline pathway (41), the final step of which involves the conversion of CDP-choline to PC. This reaction is catalyzed by CPT1 and CEPT1, the former of which localizes to the Golgi (42).

The Golgi occupies only a small volume of the whole cell (43), thus we hypothesized that Golgi-IP coupled with LC/MS can provide a powerful tool to detect subcellular metabolic changes that are missed in whole-cell measurements. To test this, we chose to analyze metabolite changes in *SLC35A2* knock-out (KO) cells, encoding a known transporter that imports UDP-hexose from the cytosol into the Golgi (44) (Fig. 4D). Because we could not detect SLC35A2 even in wildtype cells using commercially available antibodies, we verified by mass spectrometry that no detectable SLC35A2 protein was observed in the KO cell GolgiTAG-IPs (Fig. S5A). We also confirmed that Golgi-IP is as efficient in *SLC35A2* KO cells as in those with wild-type genotype (Fig. S5B). Targeted metabolic analysis revealed that UDP-hexose level was significantly reduced in Golgi derived from *SLC35A2* KO cells while its level in the whole cell was unchanged compared to wild-type cells (Fig. 4E). Importantly, the level of Golgi-associated UDP-hexose was rescued by re-expressing wild-type SLC35A2 in *SLC35A2* KO cells (Fig. 4E, Fig. S5C, S5D). These results provide *in vivo* evidence for the role of SLC35A2 in transporting UDP-hexose into the Golgi lumen. In addition to UDP-hexose, we surprisingly observed a similar level of reduction for UDP-N-acetyl-hexosamine in the Golgi upon *SLC35A2* ablation, suggesting that SLC35A2 may be able to transport both substrates into the Golgi or that both species are metabolically coupled (Fig. S5E). To further exclude the possibility that the biosynthesis of UDP-N-acetyl-hexosamine was inhibited due to the cytosolic accumulation of UDP-hexose upon SLC35A2 ablation, we incubated HEK293T whole-cell lysates with uniformly labeled U-^13^C_6_-glucose with or without unlabeled UDP-hexose supplementation. The amount of de novo synthesized U-^13^C_6_-UDP-N-acetyl-hexosamine was not decreased by UDP-hexose supplementation (Fig. S5F).

Taken together, our data show that Golgi-IP metabolomics faithfully uncovers the Golgi metabolome and identifies UDP-hexose and its derivatives as the top Golgi metabolites. In addition, Golgi-IP provides a powerful tool to study Golgi transporters *in vivo* and uncover subcellular metabolite changes that are difficult to study using whole-cell approaches.

### Lipidomic analysis of the Golgi determines its lipid content

Membrane-bound organelles have a distinct lipid composition that supports their functions (45, 46). To leverage our Golgi-IP to determine the lipid composition of the human Golgi, we performed untargeted lipidomic analysis on GolgiTAG IPs. Unbiased analysis indicated an overall strong enrichment pattern in IPs from GolgiTAG cells compared to those derived from ControlTAG ones (Fig. 5A and Table S5). We compared top-50 enriched species in GolgiTAG IPs with those that showed no enrichment, which we define as non-Golgi lipids (at the bottom of the volcano plot) (Fig. 5B). The most Golgi-enriched lipids belong to phospholipids mainly phosphatidylcholine (PC) followed by phosphatidylinositol (PI) and phosphatidylserine (PS). This finding of PCs and PIs as the most Golgi-enriched lipids is consistent with the reported roles of these lipids in coat protein complex I-mediated retrograde Golgi transport (47). Additionally, reduction of PCs impairs Golgi protein transport (48). To further characterize Golgi-enriched lipids, we also calculated the degree of unsaturation and found that Golgi-enriched lipids are slightly more unsaturated, although the difference was not statistically significant (Fig. 5C). In addition to phospholipids, we also observed enrichment of glycosphingolipids including hexosylceramides, dihexosylceramides and monosialodihexosylgangliosides (Fig. 5D). This is consistent with the role of the Golgi in catalyzing the synthesis of glycosphingolipids (49).

**Figure 5.**
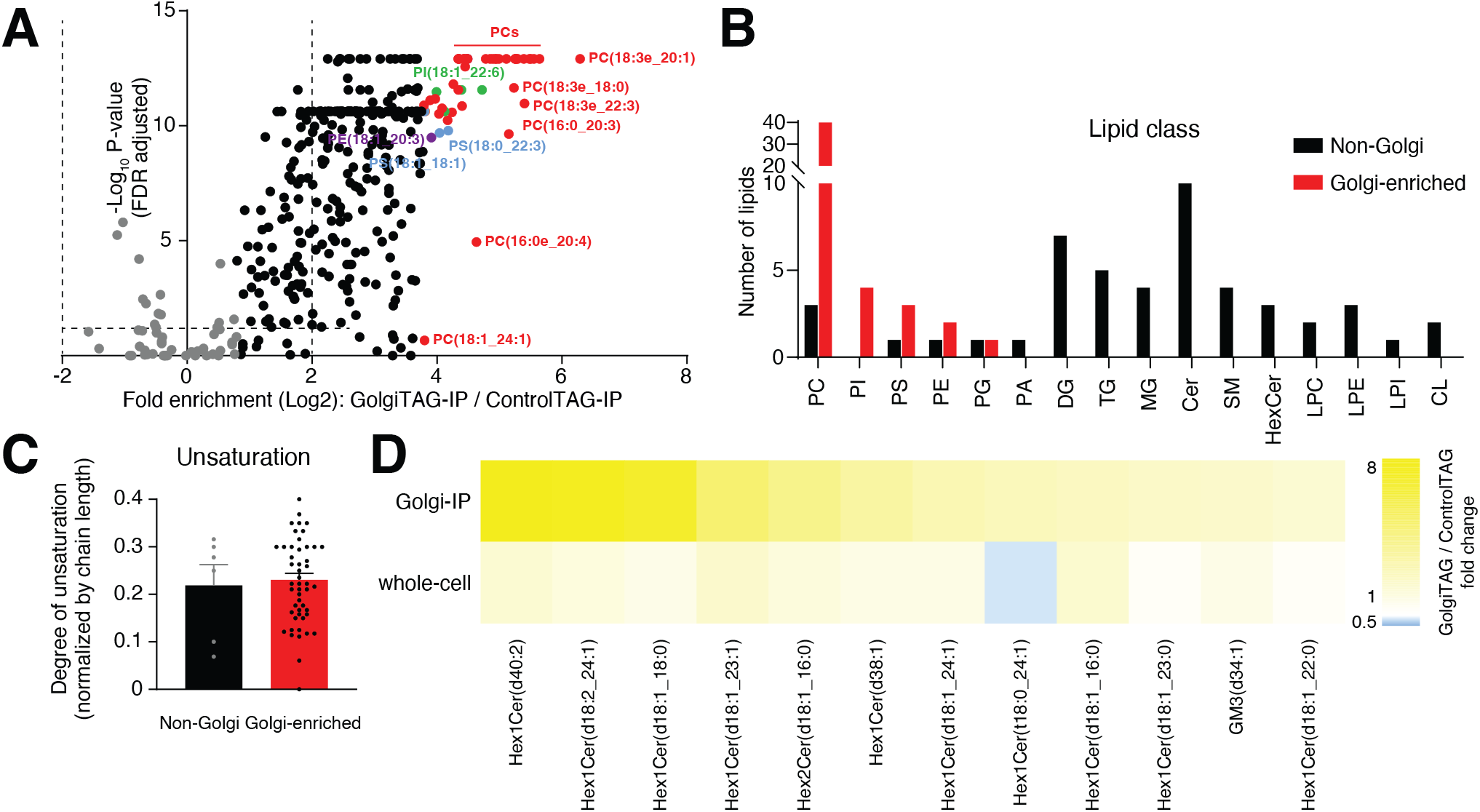
Golgi-IP lipidomics identified phospholipid as the primary Golgi lipid class. (A) Volcano plot of the fold enrichment of lipids in IPs derived from GolgiTAG compared to those from Control HEK293 cells (p-value corrected by the Benhamini-Hochberg method, FDR=5%, n= 6). Top-50 enriched species are highlighted in multi-colors. Non-Golgi lipids are in Gray. Data are in Table S5. (B) Profiling of lipid class in top-50 enriched species in the Golgi against the non-Golgi counterparts. (C) Comparison of the degree of unsaturation for phosphatides between the same groups. (D) Heatmap showing the fold change of glycolipid abundances in Golgi and whole-cell fractions in GolgiTAG and Control HEK293 cells. Full lipid names are shown in Table S5.

In summary, our results show that Golgi-IP can be used to determine the lipid composition of the Golgi and will facilitate future studies to probe its lipid content under various nutrient and disease states.

## Discussion and conclusions

Subcellular compartments provide optimal environments for specialized biochemical reactions (50). Of these compartments, the Golgi, represents a major site for protein maturation and complex lipid biosynthesis. We developed the Golgi-IP method to enable the rapid immunopurification of the intact Golgi. Our method successfully enriches for known Golgi proteins with minimal contamination from other subcellular compartments. Moreover, the Golgi-IP coupled with high-resolution mass spectrometry can be used to perform multimodal analyses of the Golgi molecular content. This has provided comprehensive maps of the human Golgi proteome, metabolome and lipidome.

Through our proteomic analysis, we confirmed previously known Golgi proteins and identified novel ones associated with this organelle. While previously annotated Golgi proteins are mostly associated with the Golgi membrane, the new candidates identified by our analysis suggest that certain proteins may be shared between the Golgi and other organelles such as the ER and lysosomes. This may be explained by the anterograde flow of the secretory pathway and the close interaction between the Golgi and these subcellular compartments (51, 52). Our results identify *SLC39A9*-encoded protein ZIP9 as a Golgi resident protein. ZIP9 is the only known zinc transporter to have a hormone receptor activity (53) and has been described to have an important role in regulating cytosolic zinc levels (54). While its Golgi-association data are limited, a previous report showed a colocalization between ZIP9 and TGN46, a *trans*-Golgi network marker (31). Our proteomic data and immunofluorescence validation support this report and establish ZIP9 as a Golgi resident protein. TM9SF1 (transmembrane 9 superfamily member 1), a protein implicated in starvation-induced autophagy, was previously shown to localize to the autophagosome and lysosome (55). We initially speculated that the enrichment of this protein in our GolgiTAG-IP was due to the proximity of the Golgi to the lysosome. However, our immunofluorescence analysis confirms that it localizes to the Golgi. KIAA2013 is an uncharacterized protein that is enriched 21-fold in our GolgiTAG-IP. Confocal imaging also confirmed it as a Golgi resident protein. These examples and others provide a proof of principle for the utility of our Golgi-IP method to study the proteomic landscape of the Golgi in health and disease states.

Through untargeted metabolomic analysis of the purified Golgi, we report the first metabolite atlas of the human Golgi. Consistent with the Golgi function in protein and lipid glycosylation, we have identified UDP-sugars and their derivatives as the top enriched polar species in the Golgi. These include UDP-containing species used for the different types of glycosylation reactions including the addition of hexose, N-acetyl-hexosamine and hexose-uronate. Furthermore, using the Golgi-IP method to analyze the metabolite content of Golgi derived from *SLC35A2* KO cells, we provided subcellular evidence that UDP-hexose is transported by SLC35A2 into the Golgi. Surprisingly, we also found a Golgi-specific reduction in the levels of UDP-N-acetyl-hexosamine, suggesting that SLC35A2 might also transport this metabolite and that UDP-hexose levels in the Golgi are metabolically coupled to those of UDP-N-acetyl-hexosamine. Of importance, none of the observed changes in the levels of these species are detected at the whole-cell level, further indicating the utility of the Golgi-IP method to study the molecular function of subcellular transporters.

Finally, using untargeted lipidomics of purified organelles, we were able to investigate the phospholipid composition of the Golgi. We observed that PCs, PIs and PSs constitute more than 90% of top-50 enriched lipids in the Golgi fraction. Consistent with our findings, PC plays a critical role in modulating Golgi protein transport to the plasma in the Chinese hamster ovary cell line MT58 (48). In addition, the enzyme PITPβ that exchanges PC and PI at the Golgi-ER interface is essential for Golgi to ER retrograde protein transport in HeLa cells (47). In the same work, it was hypothesized that PITPβ is required for importing ER-synthesized PI in exchange for PC in the Golgi, so that the level of PI4P can be maintained for proper Golgi functioning (47). We believe that Golgi-IP holds a great potential in addressing questions related to this biology and exploring the broader area of lipid metabolism in the Golgi.

While insights into organelle biology and metabolism can be achieved through different means including the combined use of multiple mass spectrometry techniques (56), customized isotope tracers (57) and fluorescence sensing (58, 59), direct isolation of organelles via immunoprecipitation has the advantage of allowing the unbiased analyses of their proteomic, metabolomic and lipidomic content (36). Additionally, the expression of the tags can be easily engineered to allow temporal, spatial and cell-type specific isolation of organelles in animal models (11, 14, 15). Therefore, we believe that our work that provides a novel technique to isolate and characterize the Golgi will advance the overarching objective to resolve compartmentalized biology with unprecedented subcellular resolution and molecular coverage.

## Materials and Methods

### Cell culture

HEK293 cells (ATCC. Cat# CRL-1573, RRID:CVCL_0045), HEK293FT (Invitrogen. Cat# R70007, RRID:CVCL_6911), HeLa (ATCC. Cat# CCL-2, RRID:CVCL_0030), HEK293T wild type and HEK293T *SLC35A2*-KO (60) were cultured in DMEM (Dulbecco’s Modified Eagle Medium, Gibco™) containing 10% (v/v) foetal calf serum, 2 mM L-glutamine, 100 U/ml penicillin and 100 μg/ml streptomycin at 37°C in a humidified incubator maintaining 5% (v/v) CO_2_. Cells were regularly tested for mycoplasma contamination. Both HEK293T wild type and HEK293T *SLC35A2* knock-out cells were gifts from Mariusz Olczak’s lab, University of Wroclaw.

### Plasmids

The plasmids (except VSVG and Gag/Pol) used in this study were obtained from the MRC PPU Reagents and Services (https://mrcppureagents.dundee.ac.uk) or other indicated source. Each plasmid was confirmed by sequencing at the MRC Sequencing and Services (https://www.dnaseq.co.uk). Plasmids are available to request via MRC PPU Reagents and Services (https://mrcppureagents.dundee.ac.uk).

| Construct | Plasmid Backbone | Source | Catalogue Number |
| --- | --- | --- | --- |
| TMEM115-3xHA | pLJC5 | MRC PPU Reagents and Services | DU68534 |
| 3xHA | pLJC5 | MRC PPU Reagents and Services | DU70022 |
| SLC35A2-2xFLAG | pBABED | MRC PPU Reagents and Services | DU72094 |
| KIAA2013-HA | pcDNA5D | MRC PPU<br>Reagents and<br>Services | DU72430 |
| TM9SF1-HA | pcDNA5D | MRC PPU<br>Reagents and<br>Services | DU72445 |
| DIPK1A-HA | pcDNA5D | MRC PPU<br>Reagents and<br>Services | DU72446 |
| C5orf22-HA | pcDNA5D | MRC PPU<br>Reagents and<br>Services | DU72447 |
| SLC39A9-HA | pcDNA5D | MRC PPU<br>Reagents and<br>Services | DU72466 |
| TMEM219-HA | pcDNA5D | MRC PPU<br>Reagents and<br>Services | DU72705 |
| ABHD13-HA | pcDNA5D | MRC PPU<br>Reagents and<br>Services | DU72706 |
| CCDC126-HA | pcDNA5D | MRC PPU<br>Reagents and<br>Services | DU72709 |
| VSVG | Lenti-X HTX | Clontech | 631247 |
| Gag/Pol | Lenti-X HTX | Clontech | 631247 |
| VSVG | pCMV | Cell Biolabs | RV-110 |
| Gag/Pol | pCMV | Cell Biolabs | RV-111 |

### Transient Transfection

Cell transfection was undertaken using the Polyethylenimine (PEI) method. Except otherwise stated, all cells were transfected at 60% confluency in a 10 cm diameter plate. Briefly, for a 10 cm diameter dish of cells, 6 μg of DNA and 18 μg of PEI were diluted in 0.5 ml of Opti-MEM™ Reduced serum medium (Gibco™), incubated for 30 minutes at room temperature, added dropwise into plates and cells incubated for 24 h at 37°C.

For immunofluorescence transient transfection analysis, 400,000 cells were seeded on ethanol-sterilised coverslip in 6 well 3.5 cm diameter plate containing 22 × 22 mm glass coverslips (VWR. Cat# 631-0125). 2 μg of DNA and 6 μg of PEI were diluted in 0.25 ml of Opti-MEM™ Reduced serum medium (Gibco™), incubated for 30 minutes at room temperature and added dropwise to plates containing cells, which were incubated for 24 h at 37°C.

### Buffers

The Triton-X100 cell lysis buffer used in this study to lyse cells and GolgiTAG immunoprecipitates contained 50 mM HEPES-KOH pH7.4, 40 mM NaCl, 2 mM EDTA, 1.5 mM NaVO_4_, 50 mM NaF, 10 mM NaPyrophosphate, 10 mM Na-β-Glycerophosphate and 1% (v/v) TritonX-100 and immediately prior to use the buffer was supplemented with cOmplete™ protease inhibitor cocktail (Roche. Cat# 04693116001).

The isotonic buffer used for homogenising cells for the GolgiTAG immunoprecipitation contains potassium-supplemented phosphate saline buffer (KPBS) (136 mM KCL, 10 mM KH_2_PO_4_. Adjust to pH7.25 with KOH).

The SDS Lysis Buffer used to lyse cells as well as the GolgiTAG immunoprecipitates for proteomics analysis, contained 100 mM HEPES-KOH pH 7.4, 2% (w/v) SDS and immediately prior to use supplemented with cOmplete™protease inhibitor cocktail (Roche. Cat# 04693116001) and PhosStop (Roche. Cat# 4906845001).

The metabolomic solubilization buffer used to resuspend whole-cell lysates and GolgiTAG immunoprecipitates was made of 80% (v/v) Optima LC/MS grade methanol (Fisher. Cat# A456-4) diluted in Optima LC/MS grade water (Fisher, Cat# W6-4). This was supplemented with 500 nM isotopically labelled amino acids (Cambridge Isotope Laboratories. Cat# MSK-A2-S).

The lipidomics solubilization buffer used to resuspend whole-cell lysates and GolgiTAG immunoprecipitates was made of LC/MS grade chloroform (VWR. Cat# 22711.260) and Optima LC/MS grade methanol (Fisher. Cat# A456-4) at a 2:1 (v/v) ratio, supplemented with Splashmix (Avanti. Cat# 330707) (1:1000 dilution).

### Cell Lysis for immunoblotting analysis

For cell lysis for immunoblotting analysis, medium is aspirated, cells placed on ice and washed twice with ice cold PBS. For a 10 cm diameter dish, 1 ml Triton-X100 lysis buffer was added, and the cells scraped into 1.5 ml tube, then incubated on ice for 10 minutes. The extract is clarified by centrifugation at 17000 x *g* for 10 minutes at 4°C. Supernatants were collected and stored in aliquots at −80°C and/or used immediately for immunoblotting.

### Immunoblotting assay

Whole cells or immunoprecipitated Golgi were lysed in 1% (v/v) Triton-X100 buffer as described above. Protein concentrations of lysates were determined using Pierce™ BCA Protein Assay Kit (Thermo. Cat# 23227) in technical duplicates according to manufacturer’s instructions. Lysates were incubated with a quarter of the volume of 4X SDS-PAGE sample buffer [50 mM Tris–HCl, pH 6.8, 2% (w/v) SDS, 10% (v/v) glycerol, 0.02% (w/v) Bromophenol Blue and 1% (v/v) 2-mercaptoethanol] (Novex). For SDS–PAGE, typically 2-20 μg samples were loaded on 4–12% Bis-tris gradient gels (Thermo Fisher Scientific. Cat# WG1402BOX or Cat# WG1403BOX) and run at 120 V for 120 minutes. Proteins were transferred onto nitrocellulose membranes at 90 V for 90 minutes on ice in transfer buffer [48 mM Tris–HCl, 39 mM glycine and 20% (v/v) methanol]. Transferred membranes were blocked with 5% (w/v) milk in TBST Buffer at room temperature for 60 minutes. Membranes were then incubated with primary antibodies diluted in 5% (w/v) BSA in TBST Buffer overnight at 4°C. After washing in TBST three times, membranes were incubated at room temperature for 1 h with near-infrared fluorescent IRDye antibodies (LI-COR) diluted 1:10,000 in TBST buffer and developed using the LI-COR Odyssey CLx Western Blot imaging system. Detailed protocol is on protocol.io (dx.doi.org/10.17504/protocols.io.bp2l61oxdvqe/v1)

### Immunofluorescence assay

The cells indicated in the figure legends were seeded on 22 x 22 mm glass coverslips in 6 well 3.5 cm diameter plates. Transfection was performed as described above for those that were transfected. Where indicated, cells were treated with 30 μM nocodazole (Sigma. Cat# M1404) generated from a 30 mM stock dissolved in DMSO 1 h prior to fixation. For fixation, medium was aspirated, and cells were fixed in 3 ml of 4% (w/v) paraformaldehyde in PBS for 10 minutes at room temperature. Fixed cells were then washed three times at 5 minutes intervals with 0.2% (w/v) bovine serum albumin (BSA) dissolved in PBS. Cells were permeabilized with 1% (v/v) Nonidet P40 diluted in PBS for 10 minutes at room temperature. Permeabilized cells were blocked with 1% (w/v) BSA in PBS for 1 h, then incubated with primary antibody diluted in PBS for 1 h at room temperature in a dark chamber. Antibody dilutions used are described in the table below. This is followed by three washes with 0.2% (w/v) BSA at 5 minutes intervals. Cells were then incubated for 1 h, in a dark chamber, with a mixture of secondary antibodies containing Alexa Fluor 594 donkey anti-Rat (Invitrogen. Cat# A21209. RRID:AB_2535795) and Alexa Fluor 488 donkey anti-rabbit (Invitrogen Cat# A21209. RRID:AB_2535792) or Alexa Fluor 488 anti-mouse (Invitrogen. Cat# A21202. RRID:AB_141607) at 1:500 dilution in PBS. Cells were washed again three times with 0.2% (w/v) BSA, rinsed in MilliQ water, and mounted on glass microscopic slides with Prolong™ Gold antifade reagent with DAPI (Invitrogen. Cat# P36931). Slides were then imaged using Leica TCS SP8 MP Multiphoton Microscope using a 40x oil immersion lens choosing the optimal imaging resolution with 1-pixel size of 63.3 nm × 63.3 nm. Detailed protocol is on protocol.io (dx.doi.org/10.17504/protocols.io.q26g74qpkgwz/v1)

### Antibodies for Immunoblotting (IB) and Immunofluorescence (IF)

| Antibody target | Company | Catalogue number (RRID) | Host species | Dilution for IB | Dilution for IF |
| --- | --- | --- | --- | --- | --- |
| ACBD3 | Sigma | HPA015594 (RRID:AB_1844491) | Rabbit | 1:1000 |  |
| ACBD3 (clone 2G2) | Sigma | WH0064746M1 (RRID:AB_2220068) | Mouse |  | 1:1000 |
| GOLGIN97 | Cell<br>Signalling<br>Technology | 13192<br>(RRID:AB_2798144) | Rabbit | 1:1000 | 1:100 |
| GM130 | Cell<br>Signalling<br>Technology | 12480<br>(RRID:AB_2797933) | Rabbit | 1:1000 | 1:3000 |
| HA | Sigma | 11867423001<br>(RRID:AB_390918) | Rat | 1:1000 | 1:1000 |
| HSP60 | Cell<br>Signalling<br>Technology | 12165<br>(RRID:AB_2636980) | Rabbit | 1:1000 |  |
| Calreticulin | Cell<br>Signalling<br>Technology | 12238<br>(RRID:AB_2688013) | Rabbit | 1:1000 |  |
| LAMP1 | Cell<br>Signalling<br>Technology | 3243<br>(RRID:AB_2134478) | Rabbit | 1:1000 |  |
| α-Tubulin | Cell<br>Signalling<br>Technology | 3873S<br>(RRID:AB_1904178) | Mouse | 1:5000 | |
| GCC185<br>(serum) | Pfeffer's lab | (35) | Rabbit |  | 1:500 |
| FLAG | Sigma | F1804<br>(RRID:AB_262044) | Mouse | 1:1000 |  |

| Secondary Antibodies | Company | Catalogue number<br>(RRID) | Dilution for IB | Dilution for IF |
| --- | --- | --- | --- | --- |
| IRDye 800CW Donkey anti-Rabbit IgG | LI-COR | 926-32213<br>(RRID:AB_621848) | 1:10,000 |  |
| IRDye 800CW Donkey anti-Mouse IgG | LI-COR | 926-32212<br>(RRID:AB_621847) | 1:10,000 |  |
| IRDye 800CW Goat anti-Rat IgG | LICOR | 926-32219<br>(RRID:AB_1850025) | 1:10,000 |  |
| Alexa Flour 594 Donkey anti-Rat | Invitrogen | A21209<br>(RRID:AB_2535795) |  | 1:500 |
| Alexa Flour 488 Donkey anti-Rabbit | Invitrogen | A21206<br>(RRID:AB_2535792) |  | 1:500 |
| Alexa Flour 488 Donkey anti-Mouse | Invitrogen | A21202<br>(RRID:AB_141607) |  | 1:500 |

### Generation of TMEM115-3xHA (GolgiTAG) stable cell lines

To stably introduce GolgiTAG into the indicated HEK293 and HEK293T cells, we used a lentivirus approach. The first part of the method involves generating lentiviral particles in HEK293FT packaging cells. For this 6 μg lentivirus cDNA construct expressing TMEM115-3xHA (GolgiTAG) was incubated with 3.8 μg pGAG/Pol (Clontech. Cat# 631247) and 2.2 μg pVSVG (Clontech. Cat# 631247) cDNAs, diluted in 0.25 ml Opti-MEM™ Reduced serum medium (Gibco™). This mixture was added to 36 μg of PEI diluted in 0.25 ml Opti-MEM™ Reduced serum medium (Gibco™). After 30 minutes of incubation at room temperature, the mixture was added dropwise into 10 cm diameter dish of 60% confluent HEK293FT cells, which were then incubated for 24 h at 37°C. 24 h post-transfection, medium was removed and replaced with 10 ml of fresh medium and left for another 24 h. This medium, containing lentivirus particles, was collected and used to generate the indicated stable HEK293 or HEK293T cells. These cells, at 60% confluence, were incubated with 5 ml medium containing lentivirus particles diluted with 5 ml fresh DMEM medium in the presence of 10 μg/ml Polybrene (Milipore. Cat# TR1003G) to promote viral infection. After 24 h the medium was replaced with a selection medium supplemented with 2 μg/ml Puromycin (Sigma. Cat# P9620). The cells that survived and proliferated in 2 μg/ml Puromycin were selected. These cells were permanently cultured in a medium containing 2 μg/ml Puromycin. A Detailed protocol for preparing these stable cell lines is described on protocol.io (dx.doi.org/10.17504/protocols.io.6qpvrdjrogmk/v1)

### Introduction of SLC35A2-2xFLAG into HEK293T SLC35A2 knock-out cells

To rescue the expression of SLC35A2 in HEK293T knock-out cells, we utilized a retrovirus approach. 6 μg retrovirus cDNA construct expressing SLC35A2-2xFLAG, was incubated with 3.8 μg pGAG/Pol (Cell Biolabs. Cat# RV-111) and 2.2 μg pVSVG (Cell Biolabs. Cat# RV-110) plasmids, diluted in 0.25 ml Opti-MEM™ Reduced serum medium (Gibco™). This mixture was added to 36 μg of PEI diluted in 0.25 ml Opti-MEM™ Reduced serum medium (Gibco™). After 30 minutes of incubation at room temperature, the mixture was added dropwise into 10 cm diameter dish of 60% confluent HEK293FT cells, which were then incubated for 24 h at 37°C. 24 h post-transfection, medium was removed and replaced with 10 ml of fresh medium and left for another 24 h. This medium, containing lentivirus particles, was collected and used to generate the indicated stable HEK293T cells. These cells, at 60% confluence, were incubated with 5 ml medium containing retrovirus particles diluted with 5 ml fresh DMEM medium in the presence of 10 μg/ml Polybrene (Milipore. Cat# TR1003G) to promote viral infection. After 24 h the medium was replaced with a selection medium supplemented with 500 μg/ml hygromycin (Invivogen. Cat# ant-hg-5). The cells that survived and proliferated in 500 μg/ml hygromycin were selected and were permanently cultured in a medium containing 500 μg/ml hygromycin. A detailed protocol is described on protocol.io (dx.doi.org/10.17504/protocols.io.kqdg3prxpl25/v1)

### Golgi-IP Method

Cells cultured on 15 cm diameter dishes were washed twice with cold PBS on ice and incubated with 1 ml ice cold KPBS. Cells were scraped on ice into a 2 ml Eppendorf tube and gently centrifuged at 1000 x *g* for 2 minutes at 4°C to pellet the cells. Supernatant is discarded and pellets are gently resuspended in 1 ml ice cold KPBS. 25 μl of the resuspended cells was taken out and used for “whole-cell fraction” analysis (see below for how this fraction is processed for different applications). The remainder of the suspension was transferred to a 2 ml glass Dounce homogeniser (VWR. Cat# 89026-386) and subjected 25 up and down cycles of homogenization. The extract was centrifuged at 1000 x *g* for 2 minutes at 4°C. The supernatant which contains cellular organelles, was collected, and transferred to a clean 1.5 ml Eppendorf tube containing 100 μl of 25% slurry of anti-HA magnetic beads (Thermo Fisher. Cat# 88837) that had been washed with KPBS Buffer. The resulting mixture was gently pipetted up and down three times and incubated on a Belly dance shaker (IBI Scientific. Cat# BDRAA115S). After incubation for 5 minutes, the tube was placed on a magnet for 30 sec and the supernatant was removed. Beads were washed 3 times with 1 ml ice cold KPBS. After the last wash, beads were resuspended with a solubilization buffer depending on downstream application. For immunoblotting studies beads were resuspended in 85 μL Triton-X100 lysis buffer for 10 minutes, beads removed, and samples centrifuged at 10, 000 x *g* for 10 minutes and supernatant transferred to a clean 1.5 ml Eppendorf tube and samples stored at −80°C. For proteomic analysis beads were resuspended with 100 μl SDS lysis buffer and incubated for 10 minutes, beads removed, and samples subjected to sonication on a Bioruptor (High intensity 30 sec-ON and 30 sec-OFF of each cycle and a total of 15 cycles were performed), and then centrifuged for 10,000 x *g* for 10 minutes. Samples were transferred to a clean 1.5 ml Eppendorf tube and stored at −80°C. For metabolomic analysis beads were resuspended in 50 μl of metabolomics solubilization buffer and left to incubate at 4°C for 10 minutes. Beads were removed and sample centrifuged for 10,000 x *g* for 10 minutes and transferred to a clean 1.5 ml Eppendorf tube and stored at −80°C. For lipidomics analysis beads are resuspended in 1 ml lipidomics solubilization buffer for 10 minutes at 4°C, beads removed, and the supernatant transferred to a clean 1.5 ml tube. The sample was incubated on a Thermomixer at 1500 rpm for 1 h at 4°C, followed by adding 200 μl 0.9% (w/v) saline (VWR. Cat# L7528) and incubation for further 10 minutes on a Thermomixer at 1500 rpm at 4°C. The mixture was centrifuged at 3000 x *g* for 10 minutes at 4°C to separate the methanol/saline (upper) and chloroform (lower) phases. The lipids are contained in the chloroform phase and the upper phase is removed and discarded. Approximately 600 μl of the lower phase was collected into a fresh 1.5 ml tube and vacuum dried on Speedvac (Thermo Scientific. Cat# SPD140DDA) before storage at −80°C.

Whole-cell fractions were processed in the following ways. For immunoblotting studies, 25 μl of cell suspension in KPBS was resuspended in 125 μl Triton-X100 lysis buffer for 10 minutes, samples centrifuged at 10, 000 x *g* for 10 minutes and supernatant transferred to a clean 1.5 ml Eppendorf tube before storage at −80°C. For proteomic analysis, 25 μl of cell suspension in KPBS was resuspended with 50 μl SDS/lysis buffer, and samples subjected to sonication on a Bioruptor (High intensity 30 sec-ON and 30 sec-OFF of each cycle and a total of 15 cycles were performed), centrifuged for 10,000 x *g* for 10 minutes. Samples were transferred to a clean 1.5 ml Eppendorf tube and stored at −80°C. For metabolomic analysis, 25 μl of cell suspension in KPBS was mixed with 225 μl of metabolomics solubilization buffer and left to incubate at 4°C for 10 minutes. Sample centrifuged at 10,000 x *g* for 10 minutes and transferred to a clean 1.5 ml Eppendorf tube and stored at −80°C. For lipidomics analysis 25 μl of extract was resuspended in 1 ml lipidomics solubilization buffer for 10 minutes on ice and the supernatant transferred to a clean 1.5 ml tube. The sample was incubated on a Thermomixer at 1500 rpm for 1 h at 4°C, followed by adding 200 μl 0.9% (w/v) saline (VWR. Cat# L7528) and incubated for another 10 minutes on a Thermomixer at 1500 rpm at 4°C. The mixture was centrifuged at 3000 x *g* for 10 minutes at 4°C to separate the methanol/saline (upper) and chloroform (lower) phases. The lipids are contained in the chloroform phase and the upper phase was removed and discarded. Approximately 600 μl of the lower phase was collected (while being careful not to take cell debris on top) into a fresh 1.5 ml tube and vacuum dried on Speedvac (Thermo Scientific. Cat# SPD140DDA) before storage at −80°C.

Detailed protocol for the GolgiTAG anti-HA immunoprecipitation step is on protocol.io (dx.doi.org/10.17504/protocols.io.6qpvrdjrogmk/v1)

### Flow cytometry analysis after GolgiTAG immunoprecipitation

Control HEK293 cells or HEK293 cells stably expressing GolgiTAG were grown in 10 cm diameter dishes to 100% confluency. The cells were treated with either DMSO or 5 μM GolgiTracker ((BODIPY™ FL C_5_-Ceramide (*N*-(4,4-Difluoro-5,7-Dimethyl-4-Bora-3a,4a-Diaza-*s*-Indacene-3-Pentanoyl)Sphingosine)) (Invitrogen. Cat# D3521) for 30 minutes at 37°C. Media were replaced with fresh media (without GolgiTracker nor DMSO) and cells were incubated for further 30 minutes. Golgi-IP was then performed, and beads washed as described above. The beads were resuspended in 500 μl isotonic buffer (KBPS) in 1.5 ml Eppendorf tube. Three replicates of 4 μl of the resuspended beads were transferred into flow cytometry tubes each containing 400 μl KPBS. Each replicate was analyzed in duplicate using a LSRFortessa™ cell analyser (BD biosciences). 50000 events were recorded and analysed in FlowJo software (RRID:SCR_008520) according to the gating strategy presented in Fig. S1A. Experiment was done in triplicates and data were statistically analyzed using One-Way ANOVA on GraphPad Prism (version 8.3.1) (RRID:SCR_002798). Detailed protocol can be found on protocols.io (dx.doi.org/10.17504/protocols.io.e6nvwk1d2vmk/v1).

### Transmission electron microscope sample preparation

Washed magnetic beads from Golgi-IP were fixed in 4% (w/v) paraformaldehyde and 2.5% (v/v) glutaraldehyde diluted in 0.1 M sodium cacodylate buffer (prepared from stock of 0.4 M sodium cacodylate buffer diluted in water and adjusted to pH7.2 with HCl) for 1 h. The beads were washed two times with 0.1 M sodium cacodylate buffer (adjusted to pH7.2 with HCl). For each wash step, beads were separated from supernatant by placing it on a magnet. Beads were then resuspended in 0.1 M sodium cacodylate buffer and centrifuged at 1000 *x g* for 1 minute, to ensure beads are tightly packed. Using a needle, the pellet was gently dislodged and a Pasteur pipette was used to transfer the pellet into a clean glass vial (VWR. Cat# 215-3571). To ensure rapid dehydration and embedding, the pellet was cut with scalpel into small pieces (about 1 mm^3^). These were then post-fixed in 1% (w/v) OsO4 with 1.5% (w/v) sodium ferricyanide diluted in 0.1M sodium cacodylate buffer for 60 minutes at room temperature. Washed three times in 0.1 M sodium cacodylate buffer (resuspend pellet in buffer, wait 1 minute for pellet to settle and use P1000 pipette to remove supernatant). This was followed by washing with water three times, then beads were incubated with 1% (w/v) tannic acid and 1% (w/v) uranyl acetate for 30 minutes at room temperature. Without further washing, pellets were gradually dehydrated through alcohol series (50%, 70%, 80%, 90%, 95% ethanol) for 10 minutes/series, then into 100% ethanol twice. The beads were then further dehydrated in 100% propylene oxide twice (10 minutes/series), then left overnight in 50% (v/v) propylene oxide and 50% (v/v) Durcupan resin (Sigma. Cat# 44611). Finally, the beads were embedded in 100% Durcupan resin. The resin was polymerised at 60°C for 48 h and sectioned on a Leica UCT ultramicrotome. Sections were contrasted with 3% (v/v) aqueous uranyl acetate and Reynolds lead citrate (1.33g lead citrate, 1.76g sodium citrate; 8 ml 1M NaOH in 50 ml water) before imaging on a JEOL 1200EX TEM using a SIS III camera. Detailed protocol is on protocol.io (dx.doi.org/10.17504/protocols.io.x54v9y9nqg3e/v1)

### Sample preparation for Quantitative proteomic analysis

The GolgiTAG immunoprecipitated bead slurry is solubilized in 100 μl of lysis buffer (2% SDS in 100mM HEPES pH8.0, Protease, and phosphatase inhibitor cocktail). The bead slurry was mixed with pipette tips a few times and allowed it to settle on the magnetic rack for 2 minutes. The supernatant was aliquoted into a new 1.5 ml Eppendorf tube and subjected to centrifugation at 17,000 *x g* for 1 minute. The samples were placed on a magnetic rack once again to remove any bead carryover and the supernatant collected into a new 1.5 ml Eppendorf tube. Samples were then subjected to sonication using Bioruptor (High intensity 30 sec-ON and 30 sec-OFF of each cycle and a total of 15 cycles were performed). In parallel, the whole cell fraction was solubilized in 50 μl of SDS lysis buffer and subjected to the Bioruptor assisted sonication and clear lysate was subjected to protein estimation using BCA assay. Both the Immunoprecipitates and whole-cell extracts were processed using S-TRAP assisted trypsin+LysC digestion as described in (61). Briefly, the proteins were reduced by adding 10 mM TCEP and incubated at 60°C for 30 minutes on a Thermo mixer at 1200 rpm agitation. The samples were then brought to room temperature and alkylated by adding 40 mM Iodoacetamide and incubated in dark at room temperature for 30 minutes on a Thermomixer at 1200 rpm agitation. Further, the samples were brought to final 5% (v/vl) SDS and added 1.2% (v/v) phosphoric acid. At this stage, six times the volume of lysate, S-Trap buffer (90% (v/v) methanol in 100mM TEABC) was added and directly loaded on S-Trap micro columns (Protifi-Co2-micro-80) and centrifuged at 1000 *x g* for 1 minute at room temperature. Columns were further washed by adding 150 μl of S-Trap buffer and centrifuged at 1000 *x g* for 1 minute and this step was repeated another three times. Finally, the S-Trap columns were transferred to new 1.5 ml Eppendorf tubes and supplemented with 1.5 μg of Trypsin+Lys-C in 70 μl and incubated at 47°C for 1.5 h on a Thermomixer followed by the temperature was set to room temperature and left overnight. The peptides were eluted by adding 40 μl of Elution buffer-1 (50mm TEABC) and then 40 μl of Elution buffer-2 (0.1% (v/v) formic acid). Further, 40 μl of Elution buffer-3 (80% (v/v) ACN in 0.1% (v/v) Formic acid) and this step was repeated two more times. The eluates were vacuum dried and solubilized in 60μl of LC-Buffer (3% ACN (v/v) in 0.1% Formic acid (v/v). The peptide amounts were measured using a nanodrop at 224 nm absorbance for an equal loading on LC-MS/MS analysis. The detailed protocol describing STRAP assisted tryptic digestion has been reported on protocols.io (dx.doi.org/10.17504/protocols.io.bs3tngnn).

### LC-MS/MS analysis for quantitative proteomics

4 μg of peptide digest was spiked with 1 μl of iRT peptides (Biognsosys). Samples were then transferred into LC glass vials. LC-MS/MS data was acquired on Orbitrap Exploris 480 mass spectrometer which is in-line with Dionex ultimate 3000 nano-liquid chromatography system. Samples were loaded onto a 2 cm pre-column (C18, 5 μm, 100 A°, 100 μ, 2 cm Nano-viper column # 164564, Thermo Scientific) at 5 μl/minute flow rate using loading pump for about 5 minutes and then resolved the peptides on a 50 cm analytical column (C18, 5 μm, 50 cm, 100 A° Easy nano spray column # ES903, Thermo Scientific) at a flow rate of 250 nl/minute flow rate by applying nonlinear gradient of solvent-B (80% (v/v) ACN in 0.1% (v/v) formic acid) for about 125 minutes with a total gradient time and run time of 145 minutes. Data were acquired in DIA-mode with a variable isolation window scheme (The isolation window scheme and key MS parameters are provided in supplemental table 1). Full MS was acquired and measured using Orbitrap mass analyzer at 120,000 resolution at m/z 200 in the mass range of 375 - 1500 m/z, AGC target was set at 300% (~ 3E6 ions) with a maximum ion injection time for 30ms. tMS2 (vDIA) scans were acquired and measured using Orbitrap mass analyzer at 30,000 resolution at 200 m/z with an AGC target of 3000% (~ 3E6 ions) with a maximum ion injection accumulation time of 70 ms. Precursor ions were fragmented using normalized higher energy collisional dissociation (HCD) using stepped collision energies of 25, 28 and 32. Both MS1 and MS2 scans were acquired in a profile mode and advanced peak determination algorithm was enabled for accurate monoisotopic envelopes and charge state determination. Loop control was set for 24 scans of tMS2 and one single MS1 scan was acquired per duty cycle. A total of 45 vDIA windows were enabled covering the mass range of 350 to 1500 m/z, the details of variableDIA isolation window values are provided in Table S3. The detailed protocol has been reported in protocols.io dx.doi.org/10.17504/protocols.io.kxygxzrokv8j/v1)

### Spectral library generation and DDA LC-MS/MS analysis

10 μg peptides from Golgi-IP were pooled and subjected to high-pH RPLC fractionation as described in (61). A total of 42 fractions were generated and each fraction was analyzed on Exploris 480 MS platform in a data dependent mode. The LC and MS instrument parameters are provided in Table S3). The detailed protocol has been reported in protocols.io dx.doi.org/10.17504/protocols.io.kxygxzrokv8j/v1)

### Proteomics database search and analysis

Pooled GolgiTAG IP DDA data were searched with MaxQuant version (1.6.10.0) (62) against a Uniprot Human database (Release July, 2020). The following search parameters were enabled: Trypsin and LysC as a protease with a maximum of two missed cleavages allowed. Deamidation of Asn and Glu, Oxidation of Met and phosphorylation Ser, Thr and Try were set as variable modifications and Carbamidomethylation of Cys was set as a fixed modification. Default search mass tolerances were used i.e. a maximum of 20 ppm MS1 tolerance and 4.5 ppm for main search and 25 ppm for MS2 was allowed. Second peptide and match between runs option was enabled. Data were filtered for 1% FDR at protein, peptide and PSM levels. The protein group file was further annotated to determine the known Golgi annotated Golgi proteins (Compartment.org and Uniprot GO terms) to map the Golgi protein abundance. The abundance rank from high to low were assigned and analysed using Perseus software version (1.6.0.15) (63). In addition, these data were also searched with the Pulsar search algorithm using Biognosys software suite to construct a hybrid library.

HEK293 cells deep proteome DDA data protein groups table was processed using Perseus software version (1.6.0.15) to estimate the protein copy numbers using Proteomic ruler method (64).

### Database search of DIA data and Data analysis

DIA datasets from GolgiTAG-IP, ControlTAG-IP and their whole-cell extracts were imported into the Spectronaut software suite (Version Rubin: 15.7.220308.50606) (65) for library free search or direct DIA to create a search archive using Pulsar search algorithm. Furthermore, this search archive was appended to the deep Golgi-tag DDA data to create a hybrid library (DDA+DIA) containing 261,484 precursors, 205,320 modified peptides and 9,629 protein groups and this library was used for the main library-based DIA search as depicted in Fig 2A The data were searched against the hybrid library and Human Uniprot database (Released July, 2021) using default Spectronaut search settings and filtered for precursor and protein Q-value cut-off of 1%. Excluded single hits and quantification were performed using MS2 area. The protein group tables were exported for further downstream analysis. The output files were further processed to map known Golgi annotated proteins, see our curated list (Table S3), known kinases and phosphatases (66, 67), and manually compiled Glycosylation, metabolism and ubiquitylation pathway components. The protein group files were further processed using Perseus software version (1.6.15.0). Missing values were imputed, and data were normalized using quantile normalization. T-test was performed between GolgiTAG-IP and ControlTAG-IP as well as between GolgiTAG-IP and GolgiTAG whole-cell extracts and between GolgiTAG whole-cell and ControlTAG whole-cell conditions and the p-values were corrected using 1% permutation-based FDR. The data visualization was further performed using in-house R scripts (Table S3) and figures were edited using Adobe illustrator version 2022.

### Untargeted metabolomics

Profiling of polar metabolites was performed on an ID-X tribrid mass spectrometer (Thermo Fisher Scientific) with an electrospray ionization (ESI) probe. A SeQuant® ZIC®-pHILIC 150 x 2.1 mm column (Millipore Sigma 1504600001) coupled with a 20 x 2.1 mm (Millipore Sigma 1504380001) guard was used to carry out hydrophilic interaction chromatography (HILIC) for metabolite separation prior to mass spectrometry. Mobile phases: A, 20 mM ammonium carbonate and 0.1% ammonium hydroxide dissolved in 100% LC/MS grade water; B, 100% LC/MS grade acetonitrile. Chromatographic gradient: linear decrease from 80-20% B from 0-20 minutes; fast linear increase from 20-80% B from 20-20.5 minutes; 80% B hold from 20.5-29.5 minutes. Flow rate, 0.15 ml/minute. Injection volume, 1.5-2.5 μL. Mass spectrometer parameters: ion transfer tube temperature, 275 °C; vaporizer temperature, 350 °C; Orbitrap resolution, 120,000; RF lens, 40%; maximum injection time, 80 ms; AGC target, 1×10^6^; positive ion voltage, 3000 V; negative ion voltage, 2500 V; Aux gas, 15 units; sheath gas, 40 units; sweep gas, 1 unit. Full scan mode with polarity switching at *m/z* 70-1000 was performed. EASYIC™ was used for internal calibration. For Data-dependent MS2 collection, pooled samples were prepared by combining replicates. HCD collision energies, 15, 30 and 45%; AGC target, 2×10^6^; Orbitrap resolution, 240,000; maximum injection time, 100 ms; isolation window, 1 m/z; intensity threshold, 2×10^4^; exclusion duration, 5 seconds; isotope exclusion, enable. Background exclusion was performed via Acquire X with one header blank and the exclusion override factor set to 3.

Compound Discoverer (Thermo Fisher Scientific) was used for initial unbiased differential analysis. In addition to online databases, we also included a local library with both masslist and mzVault spectral archives. Mass tolerance for untargeted discovery, 10 ppm; minimum and maximum precursor mass, 0-5,000 Da; retention time limit, 0-20 min; Peak filter signal to noise ratio, 1.5; retention time alignment maximum shift, 0.5 min; minimum peak intensity, 10,000; compound detection signal to noise ratio, 3. Isotope and adduct settings were kept at default values. Gap filling and background filtering were performed by default settings. Area normalization was performed by constant median after blank exclusion. Compound annotation priority: #1, MassList Search; #2, mzVault Search; #3, mzCloud Search; #4, Predicted Compositions; #5, Chemspider Search; #6, Metabolika Search. The MassList Search was customized with 5 ppm mass tolerance and 1 minute retention time tolerance. The mzVault Search was customized with 10 ppm precursor and fragment mass tolerance and 1 minute retention time tolerance. The mzCloud Search was customized with 10 ppm precursor and fragment mass tolerance. The other searches were performed with default parameters specified in the default workflow “Untargeted Metabolomics with Statistics Detect Unknowns with ID using Online Databases and mzLogic” provided by Compound Discoverer. This untargeted workflow resulted in a total of 3,356 features after the default background exclusion filter. These 3,356 features were further filtered by the following algorithm: 1) MS2 fragmentation spectra were obtained, and 2) at least 1 annotation match in the mzVault, mzCloud or Chemspider Search. To further improve the rigor of our discovery workflow, we performed additional manual filtering based on the following criteria: 1), features with retention time earlier than 3 minutes on this HILIC column, which are nonpolar and should be quantified by a C18 column, were removed, 2) features with predicted compositions containing chemical elements rarely found in human metabolome (e.g. certain halogens) were removed, and 3) features enriched in the Golgi from only one independent experiment were removed. The final filtered list contains 91 compounds shown in Table S4. Rigorous quantification of metabolite abundance was performed by TraceFinder (Thermo Fisher Scientific) in conjunction with an in-house library of known metabolite standards (MSMLS, Sigma-Aldrich). Isotopically labelled amino acids were used as internal standards. Mass tolerance for extracting ion chromatograms, 5 ppm. A detailed protocol is described on protocol.io (dx.doi.org/10.17504/protocols.io.36wgqj3p3vk5/v1 and dx.doi.org/10.17504/protocols.io.n2bvj83exgk5/v1).

### Untargeted lipidomics

Profiling of nonpolar lipids was performed on an ID-X tribrid mass spectrometer (Thermo Fisher Scientific) with a heated electrospray ionization (HESI) probe. An Ascentis Express C18 150 x 2.1 mm column (Millipore Sigma 53825-U) coupled with a 5 x 2.1 mm guard (Sigma-Aldrich 53500-U) was used to carry out C18-based lipid separation prior to mass spectrometry. Mobile phases: A, 10 mM ammonium formate and 0.1% formic acid dissolved in 60% and 40% LC/MS grade water and acetonitrile, respectively; B, 10 mM ammonium formate and 0.1% formic acid dissolved in 90% and 10% LC/MS grade 2-propanol and acetonitrile, respectively. Chromatographic gradient: isocratic elution at 32% B from 0-1.5 minutes; linear increase from 32-45% B from 1.5-4 minutes; linear increase from 45-52% B from 4-5 minutes; linear increase from 52-58% B from 5-8 minutes; linear increase from 58-66% B from 8-11 minutes; linear increase from 66-70% B from 11-14 minutes; linear increase from 70-75% B from 14-18 minutes; linear increase from 75-97% B from 18-21 minutes; hold at 97% B from 21-35 minutes; linear decrease from 97-32% B from 35-35.1 minutes; hold at 32% B from 35.1-40 minutes. Flow rate, 0.26 ml/minutes. Injection volume, 2-4 μL. Column temperature, 55 °C. Mass spectrometer parameters: ion transfer tube temperature, 300 °C; vaporizer temperature, 375 °C; Orbitrap resolution MS1, 120,000, MS2, 30,000; RF lens, 40%; maximum injection time MS1, 50 ms, MS2, 54 ms; AGC target MS1, 4×10^5^, MS2, 5×10^4^; positive ion voltage, 3250 V; negative ion voltage, 3000 V; Aux gas, 10 units; sheath gas, 40 units; sweep gas, 1 unit. HCD fragmentation, stepped 15%, 25%, 35%; data-dependent tandem mass spectrometry (ddMS2) cycle time, 1.5 s; isolation window, 1 m/z; microscans, 1 unit; intensity threshold, 1.0e4; dynamic exclusion time, 2.5 s; isotope exclusion, enable. Full scan mode with ddMS2 at *m/z* 250-1500 was performed. EASYIC™ was used for internal calibration. LipidSearch and Compound Discoverer (Thermo Fisher Scientific) were used for unbiased differential analysis. Lipid annotation was acquired from LipidSearch with the precursor tolerance at 5 ppm and product tolerance at 8 ppm. The mass list is then exported and used in Compound Discoverer for improved alignment and quantitation. Mass tolerance, 10 ppm; minimum and maximum precursor mass, 0-5,000 Da; retention time limit, 0.1-30 min; Peak filter signal to noise ratio, 1.5; retention time alignment maximum shift, 1 min; minimum peak intensity, 10,000; compound detection signal to noise ratio, 3. Isotope and adduct settings were kept at default values. Gap filling and background filtering were performed by default settings. The MassList Search was customized with 5 ppm mass tolerance and 1 minute retention time tolerance. Area normalization was performed by constant median after blank exclusion. A detailed protocol is described on protocol.io (dx.doi.org/10.17504/protocols.io.5qpvor3dbv4o/v1 and dx.doi.org/10.17504/protocols.io.3byl4jq6jlo5/v1).

Degree of saturation for phosphatides was normalized by chain length based on the following equation:

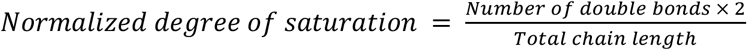

### Data Availability

Proteomics MS raw data for DDA (Spectral library) and DIA analysis was submitted to ProteomeXchange PRIDE repository (68), the data can be accessed using the dataset identifier, PXD038046. R and Python scripts associated with the Proteomics data have been deposited on the Zenodo data repository (10.5281/zenodo.7347506). Raw files for immunoblot, Confocal Immunofluorescence imaging, flow cytometry and Transmission Electron micrograph, have been deposited on the Zenodo data repository (10.5281/zenodo.7014764). All plasmids generated at the MRC Protein Phosphorylation and Ubiquitylation Unit at the University of Dundee can be requested through our website https://mrcppureagents.dundee.ac.uk/. Raw MS data files for metabolomics and lipidomics have been submitted to MetaboLights (identifier number MTBLS6511, URL: www.ebi.ac.uk/metabolights/MTBLS6511) (69)

## Author Contributions

Conceptualization, M.A.-R., D.R.A., R.F., W.D. and R.N.; Methodology, R.F., W.D., R.N., D.R.A. and M.A-R.; Experiments, R.F., W.D., R.N., E.S.R., M.I., K.N., T.K.P, E.B., and A.P.; Proteomic analysis, R.N. T.K.P; Metabolic analysis, W.D.; Manuscript writing, R.F., W.D. and R.N. wrote the manuscript and M.A.-R. and D.R.A. edited it; Funding Acquisition, M.A.-R. and D.R.A.

## Acknowledgements

The authors would like to thank the members of the Abu-Remaileh and Alessi Labs for helpful insights. We also thank the group of Mariusz Olczak from the University of Wroclaw for kindly providing us with the SLC35A2 knock-out and SLC35A2 wildtype HEK293T cells and Suzanne Pfeffer for discussion, advice and providing us with anti-GCC185 antibody. We also thank the excellent technical support of the MRC protein phosphorylation and ubiquitylation unit (PPU) DNA sequencing service (coordinated by Gary Hunter), the tissue culture team (coordinated by Edwin Allen), cloning team (coordinated by Dr Rachel Toth) antibody and protein purification team (coordinated by Dr James Hastie) and mass spectrometry team (coordinated by Dr Renata Filipe Soares). We also thank the Metabolomics knowledge center at Stanford ChEM-H (directed by Dr. Yuqin Dai). This research was funded in whole or in part by Aligning Science Across Parkinson’s [ASAP-000463] through the Michael J. Fox Foundation for Parkinson’s Research (MJFF) to M.A.-R. and D.R.A. For the purpose of open access, the author has applied a CC-BY public copyright licence to all Author Accepted Manuscripts arising from this submission. M.A.-R. is supported by NIH (DP2-CA271386) and Stanford Alzheimer’s Disease Research Center (ADRC). ER was supported by Stanford Chemical Engineering Research Experience for Undergraduates Program. M.A.-R. is a Terman Fellow and Pew-Stewart Scholar.

---

**Supplementary Figure 1:**
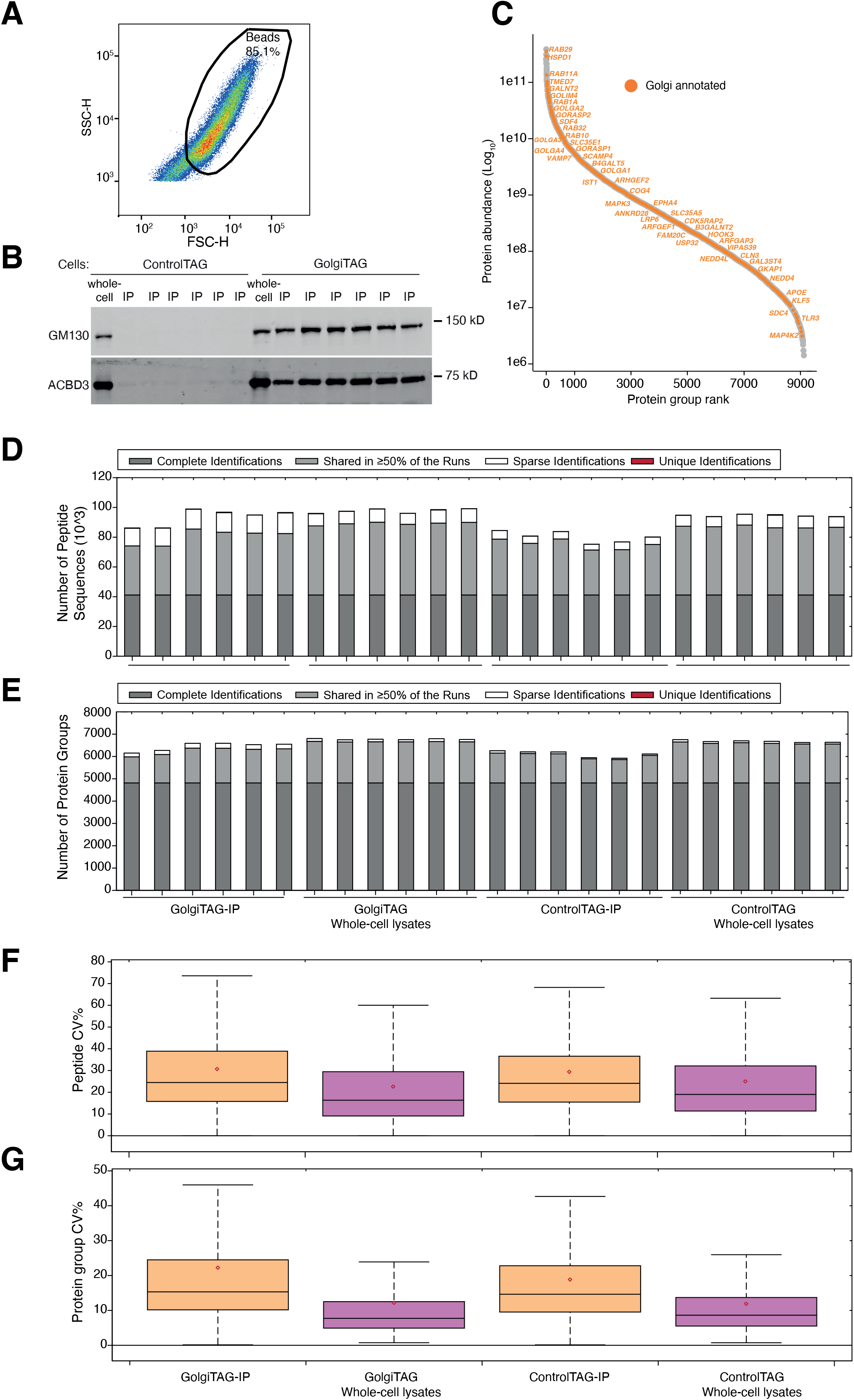
Further analysis of the GolgiTAG immunoprecipitates. (A) The gating strategy for the beads bound to Golgi for experiment presented in Fig. 1F. The beads bound to Golgi were first gated and selected as indicated, based on complexity (side scatter SSC, Y-axis) and bead size (forward side scatter FSC, X-axis). (B) Immunoblot analyses of the samples used in Fig. 2B. The whole cell lysates (2 μg) as well as the resuspended GolgiTAG immunoprecipitates (2 μg) were subjected to immunoblotting with the indicated antibodies that are markers for the Golgi. (C) Deep proteomic profiling of pooled GolgiTAG and control immunoprecipitations. Peptides were subjected to bRPLC fractionation, MS data were acquired in DDA mode to generate a spectral library which was further used for DIA data search. Rank abundance plot depicting the protein log intensities on y-axis and protein group abundance rank on x-axis. Known Golgi annotated proteins highlighted in filled circles in orange colour. (D-E) The peptide and protein groups identification summary in each sample from DIA-MS proteomic analysis of GolgiTAG-IP, ControlTAG-IP, and whole-cell lysates (n=6 in each category). (F-G) Box plots depicting the peptide and protein groups coefficient of variation from DIA-MS proteomic analysis of GolgiTAG-IP, ControlTAG-IP, and whole-cell lysates (n=6 in each category).

**Supplementary Figure 2:**
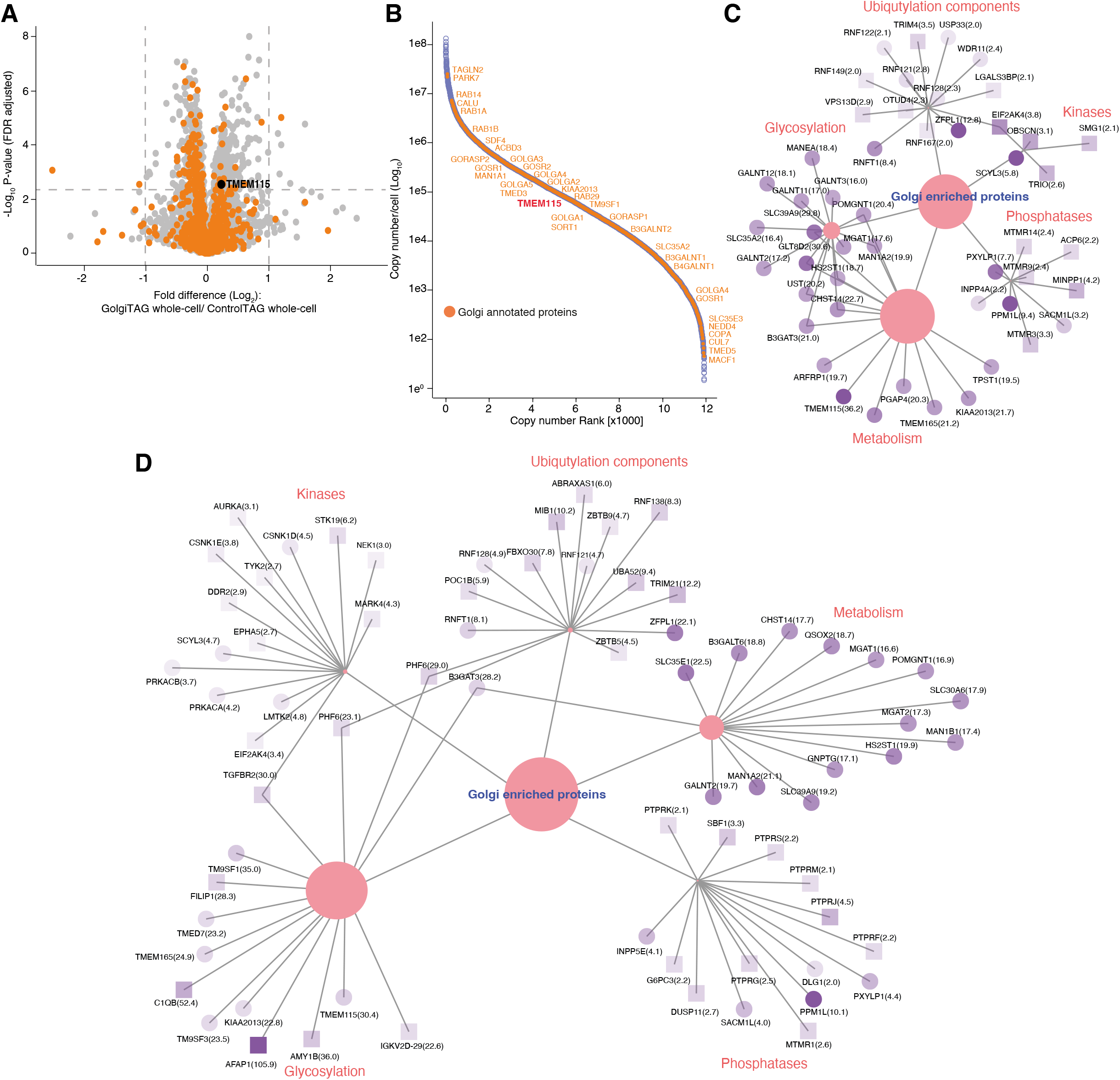
Further proteomic analysis of the GolgiTAG immunoprecipitates. (A) Volcano plot showing the fold difference between the whole-cell lysates from GolgiTAG and ControlTAG HEK293 cells (p-value adjusted for 1% Permutation-based FDR correction, s0=0.1). The orange dots depict known Golgi annotated proteins curated from databases described in (Table S3). (B) Rank abundance plot depicting the protein copy numbers from whole-cell extracts of HEK293 cells. X-axis representing the protein abundance rank (High-Low) and Y-axis representing the estimated copy numbers in log scale. The curated Golgi proteins are shown in filled circles in orange color along with the gene names for selected Golgi proteins (Table S3). (C) Network analysis for enriched proteins in GolgiTAG over control IPs. The proteins enriched in the the GolgiTag IPs versus the control (Table S1) were subjected to cytoscape network analysis ((70) using a custom python script (Table S3) and filtered for a fold change value of > 2.0 and 1% FDR for p-value significance. We have selected enzymes involved ubiquitylation (ligases and deubiqutylases), phosphorylation (kinases and phosphatases) and glycosylation. Proteins previously annotated as Golgi are shown in circles and proteins not previously annotated as Golgi shown in squares. The fold change values are represented as dark (high fold change) to light (low fold) purple color. (D) same as (C) except for those enriched by comparing the GolgiTAG IP to the wholecell lysates using data from Table S1.

**Supplementary Figure 3:**
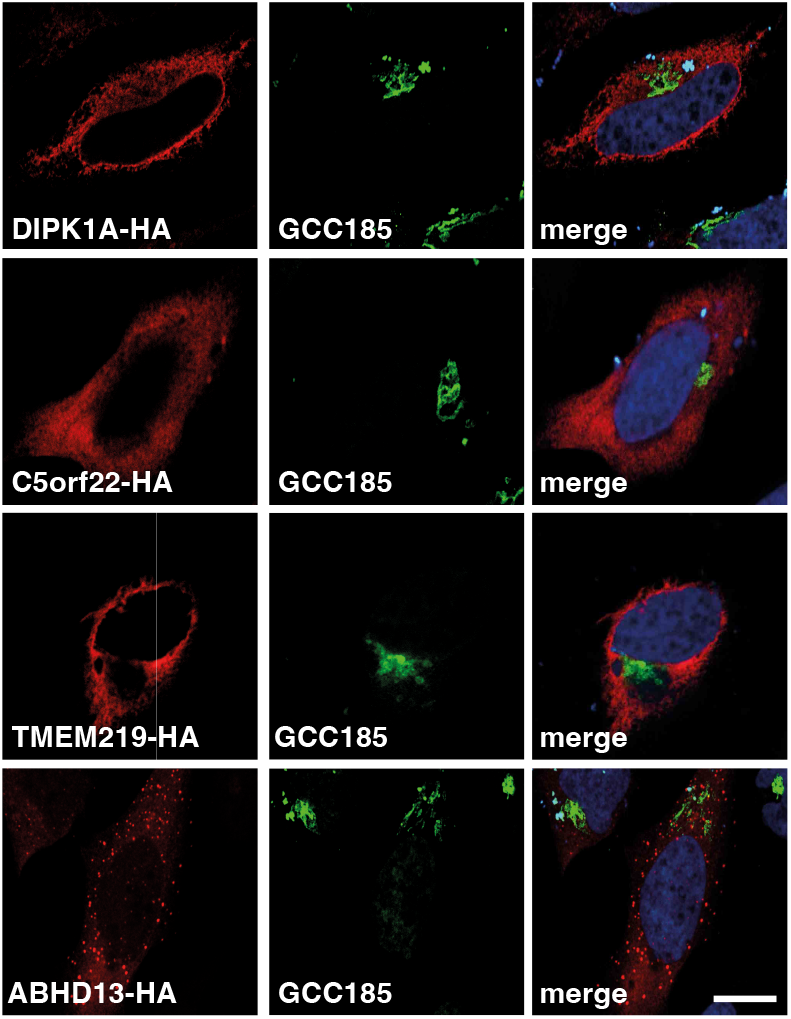
Immunofluorescence analyses of enriched proteins from GolgiTAG IPs that were not previously Golgi annotated. HeLa cells were transiently transfected with plasmids encoding for the expression of the indicated proteins fused to a C-terminal HA tag. 24h post transfection cells were the fixed with 4% (w/v) paraformaldehyde at room temperature and stained with rat anti-HA (red, left panels) and rabbit anti-GCC185 (green, middle panels). The Right panel displays the merged red and green channel. Nuclei were stained with DAPI (blue). Scale bar is 1 μm.

**Supplementary Figure 4:**
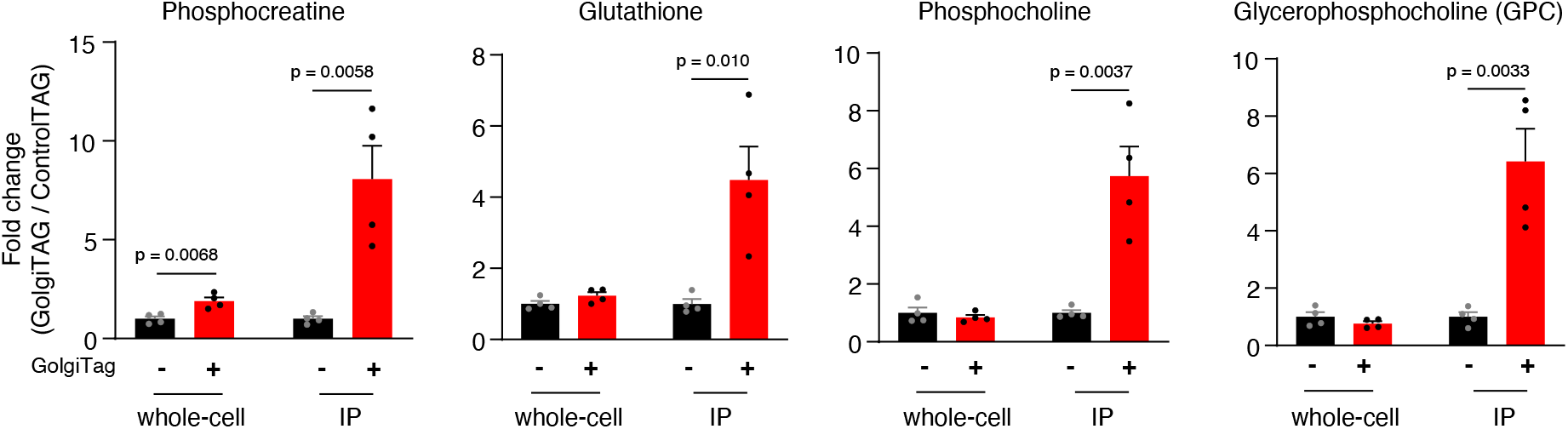
Quantitation of selected Golgi-enriched metabolites. Quantitation of phosphocreatine, glutathione, phosphocholine and glycerophosphocholine (GPC). Fold changes in the abundance of the indicated metabolites in IP and whole-cell fractions of GolgiTAG compared to those of ControlTAG cells are presented as mean ± SEM (n= 4).

**Supplementary Figure 5:**
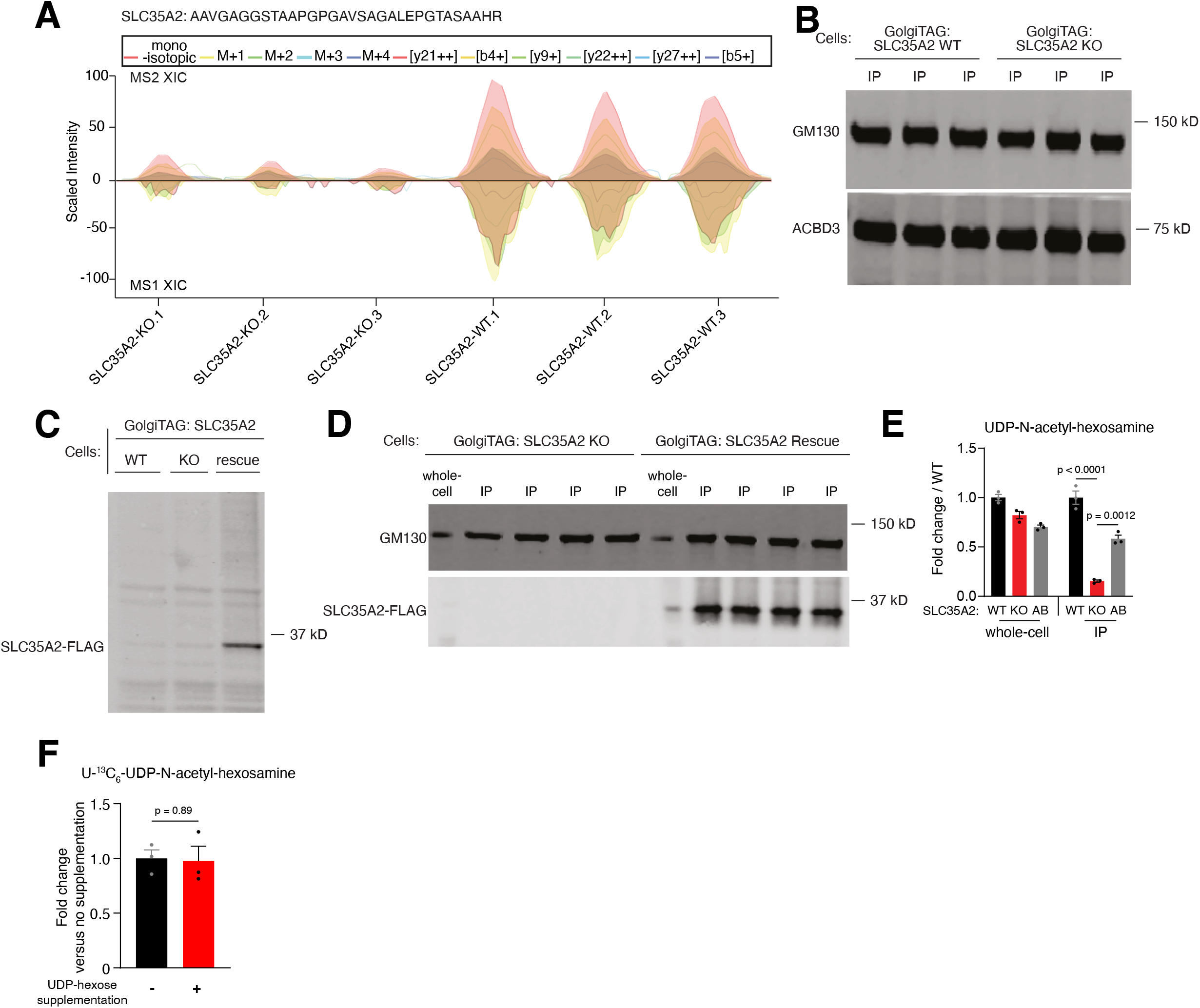
Analyzing *SLC35A2* wild type, knockout and reconstituted HEK293T cell lines. (A to B) Golgi-IP in *SLC35A2* wild type and knockout HEK293T cells validate the loss of SLC35A2 protein in the knockout cells. (A) The GolgiTAG immunoprecipitates were analyzed by DIA mass spectrometry and the extracted ion chromatography (XIC) of an SLC35A2 selective peptide (AAVGAGGSCAAPGPGAVSAGALEPGTASAAHR) which displayed robust intensity in wild type cells while lost in the knockout cells is presented. (B) Immunoblot analyses of the GolgiTAG IP from the indicated cell lines. 2 μg lysates were subjected to immunoblotting with the indicated antibodies that are markers for the Golgi. (C) Wild type, *SLC35A2* knockout and *SLC35A2* knockout HEK293T cells reconstituted with wild type SCL35A2 fused to the C-terminal Flag tag (rescue) all stably expressing the GolgiTAG were lysed in a buffer containing 1% (v/v) Triton-X100 and subjected to immunoblot analysis with the Flag antibody. (D) SLC35A2 knockout and rescued HEK293T cells were subjected to Golgi-IP. The whole cell lysates (2 μg) as well as the resuspended GolgiTAG IP (2 μg) were analyzed by immunoblotting with the indicated antibodies. (E) Targeted analysis showing fold change in the abundance of UDP-N-acetyl-hexosamine in the IP and whole-cell fractions of *SLC35A2* KO and rescued (AB) HEK293T cells compared to those from WT cells. Data are presented as mean ± SEM (n= 3). Statistical tests: two-tailed unpaired t-test. (F) Quantitation of uniformly labeled U-13C6-UDP-N-acetyl-hexosamine with or without supplementation of unlabeled UDP-hexose in whole-cell lysates from HEK293T cells. Data are presented as mean ± SEM (n= 3).

## Supplementary Tables

**Supplementary Table 1:** DIA-based quantitative proteomic analysis of immunoprecipitates and whole-cell lysates derived from GolgiTAG and ControlTAG HEK293 cells.

**Supplementary Table 2:** Pooled GolgiTAG-IP Data dependent Acquisition (DDA)-based MS analysis used to generate Spectral library for DIA.

**Supplementary Table 3:** List of manually curated Golgi proteins list.

**Supplementary Table 4:** Untargeted metabolomics of Golgi derived from GolgiTAG and ControlTAG HEK293 cells.

**Supplementary Table 5:** Untargeted lipidomics of Golgi derived from GolgiTAG and ControlTAG HEK293 cells.

